# Targeting NAAA counters dopamine neuron loss and symptom progression in mouse models of Parkinson’s disease

**DOI:** 10.1101/2022.02.10.479850

**Authors:** Francesca Palese, Silvia Pontis, Natalia Realini, Alexa Torrens, Faizy Ahmed, Francesca Assogna, Clelia Pellicano, Paola Bossù, Gianfranco Spalletta, Kim Green, Daniele Piomelli

## Abstract

The lysosomal cysteine hydrolase *N*-acylethanolamine acid amidase (NAAA) deactivates the lipid-derived mediator palmitoylethanolamide (PEA), an endogenous PPAR-α agonist that is critically involved in the control of inflammation and nociception. In this study, we asked whether NAAA-regulated PEA signaling might contribute to the pathogenesis of Parkinson’s disease (PD), a neurodegenerative disorder characterized by progressive loss of nigrostriatal dopamine neurons. Analyses of postmortem brain cortex and premortem blood-derived exosomes found elevated levels of NAAA expression in persons with PD compared to age-matched controls. Furthermore, in vitro experiments showed that the dopaminergic neurotoxins, 6-hydroxydopamine (6-OHDA) and 1-methyl-4-phenyl-1,2,3,6-tetrahydropyridine (MPTP), enhanced NAAA expression and lowered PEA content in human SH-SY5Y cells. A similar effect was observed in dopamine neurons and, subsequently, in microglia following 6-OHDA injection in mice. Importantly, deletion of the *Naaa* gene or pharmacological inhibition of NAAA activity markedly attenuated both dopamine neuron death and parkinsonian symptoms in mice treated with 6-OHDA or MPTP. The results identify NAAA-regulated PEA signaling as a control node for dopaminergic neuron survival and a potential target for therapeutic intervention in PD.

## INTRODUCTION

Parkinson’s disease (PD) is a neurodegenerative disorder characterized by the progressive loss of nigrostriatal dopaminergic neurons of the substantia nigra (SN) and their axonal projections to the basal ganglia as well as by the accumulation of α-synuclein in Lewy bodies (Bloem *et al*, 2021; Vazquez-Velez & Zoghbi, 2021; Surmeier, 2018). The underlying causes of neuronal degeneration in PD are unknown, but presumably involve both cell-intrinsic and cell-extrinsic factors, including impaired intracellular trafficking to lysosomes (Udayar *et al*, 2022; Menzies *et al*, 2015) and microglia-driven neuroinflammation (Borst *et al*, 2021). Current therapies for PD only treat disease symptoms, thus an urgent goal for research is the discovery of control nodes that may be targeted to delay or halt the loss of midbrain dopamine neurons and its neurological and psychiatric consequences (Vijiaratnam *et al*, 2021).

The lysosomal cysteine hydrolase, *N*-acylethanolamine acid amidase (NAAA) (Ueda *et al*, 2010; Piomelli *et al*, 2020), deactivates the lipid-derived mediator palmitoylethanolamide (PEA), an endogenous agonist of the ligand-operated transcription factor peroxisome proliferator-activated receptor-α (PPAR-α) (Pontis *et al*, 2016). In peripheral organs, NAAA-regulated PEA signaling at PPAR -α modulates inflammatory and nociceptive responses (Piomelli *et al*, 2020; Piomelli & Sasso, 2014) while, in the spinal cord, it counters autoimmune neuroinflammation (Migliore *et al*, 2016; Pontis *et al*, 2020; Sgroi *et al*, 2021) and controls pain chronification after end-organ injury (Fotio *et al*, 2021b). Thus far, investigations on the functions served by this signaling complex at higher levels of the neuraxis have focused on dopamine signaling (Sagheddu *et al*, 2020) and substance use (Sagheddu *et al*, 2019; Fotio *et al*, 2021a). Nevertheless, two large-scale studies have also pointed to a possible role for NAAA in neurodegenerative disorders: an analysis of three independent cohorts of persons with late-onset Alzheimer’s disease uncovered a strong association between neuronal loss and elevated *Naaa* transcription (Readhead *et al*, 2018), while an unbiased gene-array survey of postmortem human brains found abnormally high levels of *Naaa* mRNA in the SN of women with sporadic PD (Papapetropoulos *et al*, 2006).

Prompted by these findings, in the present study we conducted a series of human and animal experiments to determine whether NAAA-regulated PEA signaling might play a role in the pathogenesis of PD. We found that NAAA expression is significantly elevated in premortem blood-derived exosomes and postmortem brain cortex of persons with PD, relative to age- and sex-matched control subjects. We further found that *(i)* the dopaminergic neurotoxins 6-hydroxydopamine (6-OHDA) and 1-methyl-4-phenyl-1,2,3,6-tetrahydropyridine (MPTP) stimulate NAAA expression in human SH-SY5Y cells as well as in mouse nigrostriatal dopaminergic neurons and microglia; and *(ii)* deletion of the *Naaa* gene or pharmacological inhibition of NAAA activity strongly attenuates dopamine neuron death and parkinsonian symptoms caused by either of the two toxins. The results identify NAAA-regulated PEA signaling as a previously unrecognized molecular checkpoint for dopaminergic neuron survival and a potential target for PD modification.

## RESULTS

### NAAA expression is abnormally elevated in persons with PD

Confirming and extending prior gene array analyses (Papapetropoulos *et al*, 2006), we found that *Naaa* mRNA and NAAA protein levels were significantly elevated in postmortem specimens of brain cortex obtained from a group of persons with PD (n=43; 32 women) relative to age-matched controls (n=46; 37 women) (Fig. 1A, B; demographic and clinical data are reported in Supplementary Table 1). Stratification by sex showed that the difference was primarily driven by female patients, who were overrepresented in both control and PD samples (Fig. 1A, B). Since postmortem conditions influence the quality of results obtained with human brain tissue (Stan *et al*, 2006), we also measured NAAA protein levels in blood-derived exosomes from a separate cohort of patients with PD (n=11; 7 women; average age at collection: 54.7 ± 14.8 years, disease stage III; mean ± SD) and age-matched control subjects (n=12; 8 women; average age: 49.2 ± 10.7 years) (demographic and clinical data: Supplementary Table 2). In this case too, we observed a marked NAAA elevation in persons with PD compared to controls, with female patients accounting for the majority of the effect (Fig. 1C, D). Immunohistochemical analyses of brain cortex from a female subject with PD showed that immunoreactive NAAA was primarily localized to neurons and microglia (Fig. 1E and F; separate channels: Supplementary Fig. 1). As previously reported for mouse spinal cord (Pontis *et al*, 2020), NAAA immunoreactivity was undetectable in human cortical astrocytes (Fig. 1G; separate channels: Supplementary Fig. 1) even though these cells contain high levels of *Naaa* mRNA compared to other cortical cell lineages (http://neuroexpresso.org). The finding that NAAA levels may be elevated in persons with PD both premortem and postmortem is suggestive of a possible contribution of this enzyme to pathogenesis.

**Figure 1:**
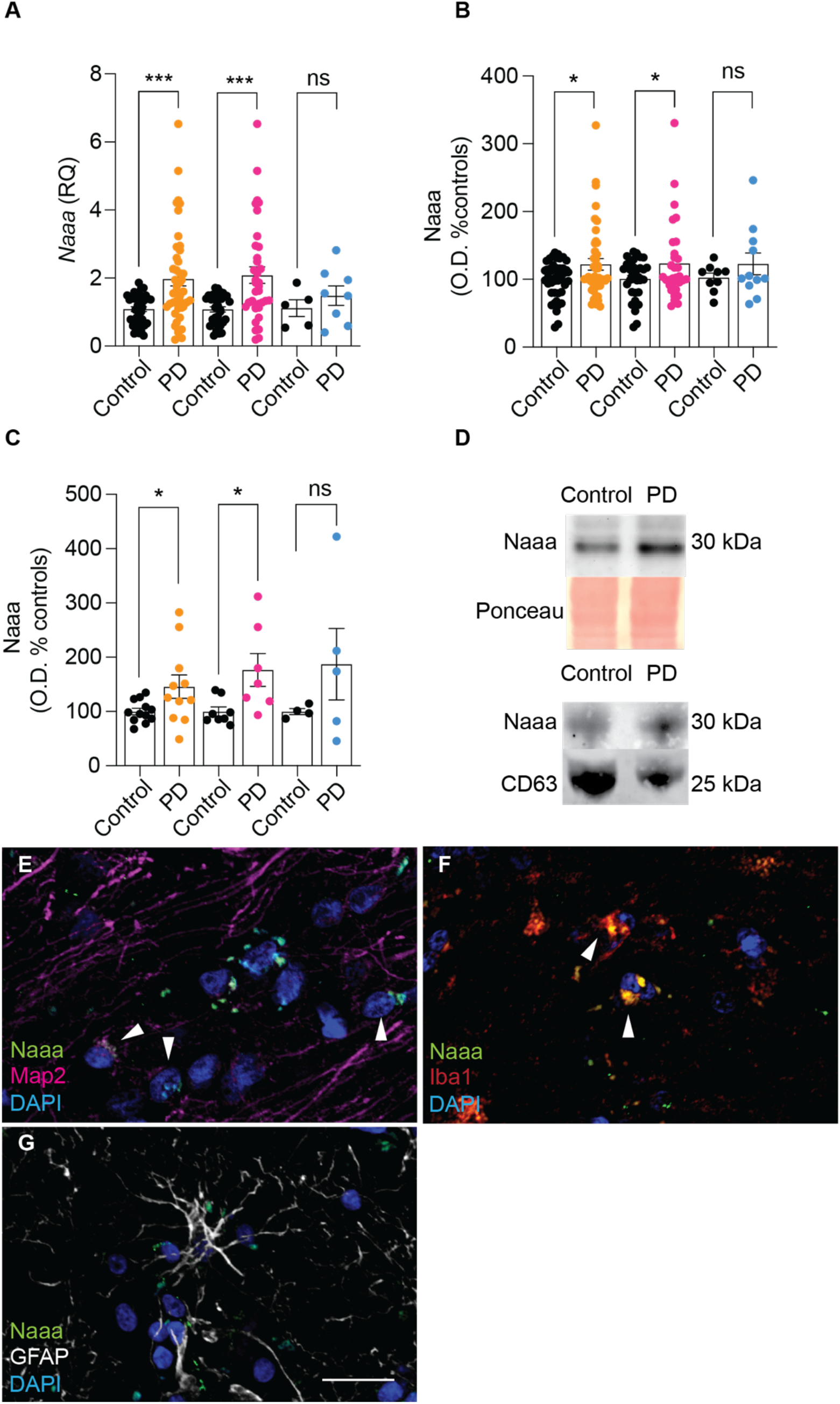
NAAA is increased in persons with PD. • (A) Quantitative RT-PCR detection of *Naaa* expression levels in postmortem cortex specimens from persons with PD (n=43) and control subjects (n=46). • (B) Densitometric analyses for NAAA protein levels in postmortem cortex specimens from persons with PD (n=43) and matched controls (n=46). • (C) Western blot analysis for NAAA protein levels in circulating exosomes obtained from plasma from persons with PD (n=11) and healthy controls (n=12). • (D) Representative blots: Ponceau staining was used for normalization of the postmortem cortex samples, CD63 for exosomes samples. • (E-G) Representative confocal pictures of immuno-staining in postmortem cortex samples from persons with PD. Merged immunofluorescence images for NAAA (green, E-G), Map2 (magenta, E), Iba1 (red, F), GFAP (white, G), cell nuclei were stained with 4’,6-diamidino-2-phenylindole (DAPI, blue). Scale bar, 20 μm. • Data information: in (A-C) Data are arranged both sexes together (orange dots) or stratified per gender: women (pink dots) and men (blue dots). ***P<0.001, *<0.05 Student’s *t*-test.

### Dopaminergic neurotoxins stimulate NAAA expression in human SH-SY5Y cells

As a first test of this idea, we examined whether two neurotoxins that cause parkinsonian symptoms in rodents and humans – 6-OHDA and 1-methyl-4-phenylpyridinium (MPP^+^, the bioactive metabolite of MPTP) (Dauer & Przedborski, 2003; Langston *et al*, 1983) – affect NAAA expression in human catecholaminergic SH-SY5Y cells (Zhao *et al*, 2017; Paliga *et al*, 2018; Urano *et al*, 2018). Incubation with 6-OHDA (100 μM) produced substantial increases in NAAA mRNA (Fig. 2A) and protein (Fig. 2B, C), which were accompanied by a decrease in cellular PEA levels (Fig. 2D). Of note, and in line with our premortem human data (Fig. 1C, D), 6-OHDA also stimulated the release of NAAA-containing exosomes into the culture medium (Fig. 2E, F). Similarly, exposing SH-SY5Y cells to MPP^+^ (2 mM) stimulated NAAA expression (Fig. 2G-I) and produced a trend toward decreased PEA content (Fig. 2J). The shared ability of 6-OHDA and MPP^+^ to induce NAAA expression in SH-SY5Y cells is consistent with an involvement in dopaminergic neuron survival. To further evaluate this possibility, we turned to in vivo mouse models.

**Figure 2.**
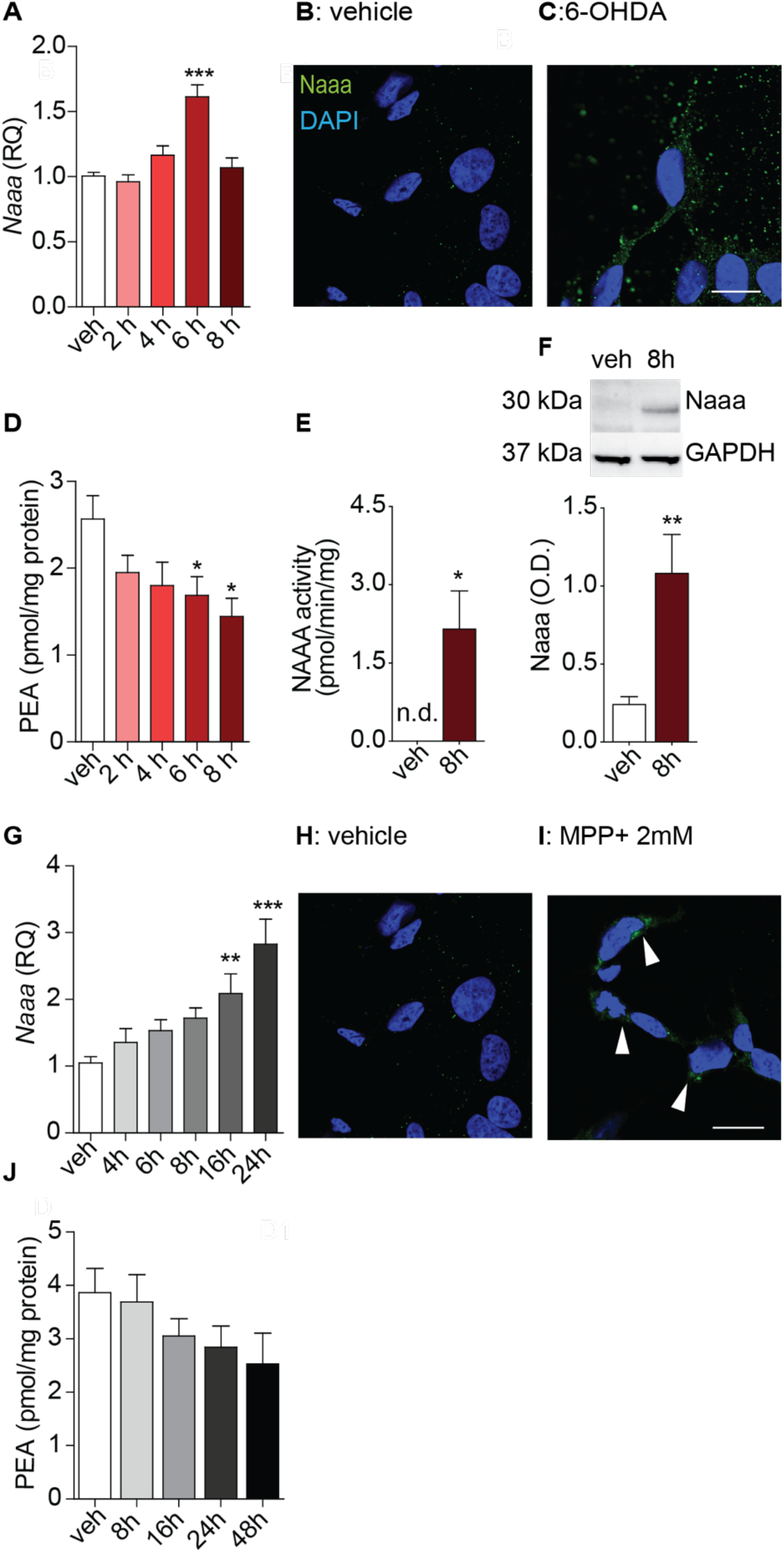
6-OHDA and MPTP induce NAAA expression in human SH-SY5Y cells. • (A) Quantitative RT-PCR detection of *Naaa* expression levels in SHSY5Y cells after incubation with 100 μM 6-OHDA (2-8h) or vehicle (veh), ***P<0.001, one-way ANOVA with Bonferroni (2 biological replicates, each analyzed in triplicate). • (B-C) Immunofluorescence images of NAAA (green) in SHSY5Y cells after incubation with vehicle (B) or 6-OHDA (C). Scale bar, 10 μm; cell nuclei were stained with DAPI (blue). • (D) PEA levels in SHSY5Y cells after incubation with 100 μM 6-OHDA (2-8h) or vehicle (veh), *P<0.05, one-way ANOVA with Bonferroni (n=6). • (E-F) NAAA activity (E) and protein quatification (F) in exosomes released by SH-SY5Y cells. **P<0.01, *P<0.05 Student’s *t*-test (n=3). • (G) Quantitative RT-PCR detection of *Naaa* expression levels in SHSY5Y cells after incubation with 2 mM MPP^+^ (2-8h) or vehicle (veh), ***P<0.001, **P<0.01 one-way ANOVA with Bonferroni (2 biological replicates, each analyzed in triplicate). • (H-I) Immunofluorescence images of NAAA (green) in SHSY5Y cells after incubation with vehicle (H) or MPP^+^ (I). Scale bar, 10 μm; cell nuclei were stained with DAPI (blue). • (J) PEA levels in SHSY5Y cells after incubation with 2 mM MPP^+^ (2-8h) or vehicle (veh), (n=6).

### 6-OHDA enhances NAAA expression in mouse nigrostriatal dopamine neurons

Unilateral striatal injections of 6-OHDA produced in male mice a rapid and persistent increase of NAAA immunoreactivity in dopaminergic tyrosine hydroxylase-positive (TH^+^) neurons of the ipsilateral SN pars compacta (Fig. 3A-D; separate channels: Supplementary Fig. 2, 3). Within 48 h of toxin administration, NAAA levels in the SN rose by approximately 30% (Fig. 3E), while the fraction of TH^+^ cells that also expressed NAAA rose from <1% to >15% and remained significantly elevated (≈6% of total) for the following 12 days (Fig. 3F; representative images and colocalization analysis: Supplementary Fig. 4). Unbiased proteomic profiling of nigral tissue at the 48-h time-point revealed that heightened NAAA expression was part of a broad reaction to the toxin that involved changes in 417 quantifiable proteins (from a total of 1164 detected; Supplementary Table 3 and Supplementary Fig. 5), most of which (378/417) were underrepresented in damaged SN (Supplementary Table 4). Consistent with the mechanism of action of 6-OHDA, which is known to impair mitochondrial complex I function (Schober, 2004; Kupsch *et al*, 2014), analysis of downregulated proteins revealed enrichment in components of oxidative phosphorylation (particularly complex I) as well as pathways of fatty-acid and glycan metabolism (Supplementary Fig. 6; Supplementary Tables 6-8). Parallel lipid analyses showed that the early surge in NAAA expression was accompanied by a ≈30% decrease in nigral PEA content (Fig. 3G), whereas levels of the less-preferred NAAA substrate, oleoylethanolamide (OEA) (Ueda *et al*, 2013; Ghidini *et al*, 2021) were only marginally affected (Fig. 3H). Two weeks after 6-OHDA injection, when inflammation had spread to the dorsolateral striatum – as evidenced by enhanced transcription of inflammatory cytokines (Fig. 4A) and damage to striatal TH^+^ fibers (Supplementary Fig. 7) -NAAA levels rose sharply in ionized calcium-binding adaptor molecule 1-positive (Iba-1^+^) cells that displayed the characteristic morphology of reactive microglia (Fig. 4 B-D). NAAA-positive microglial cells were primarily found in proximity of damaged nigral TH^+^ neurons (Fig. 3C, D; separate channels: Supplementary Fig. 3) and striatal TH^+^ fibers (Fig. 4E, F; separate channels: Supplementary Fig. 7). The results are consistent with those previously obtained in human SH-SY5Y cells (Fig. 2A-D) and indicate that enhanced NAAA expression in midbrain dopamine neurons may be an early component of the toxic response to 6-OHDA in mice. As neuroinflammation develops, NAAA induction extends to reactive microglia mobilized to damaged neural structures.

**Figure 3.**
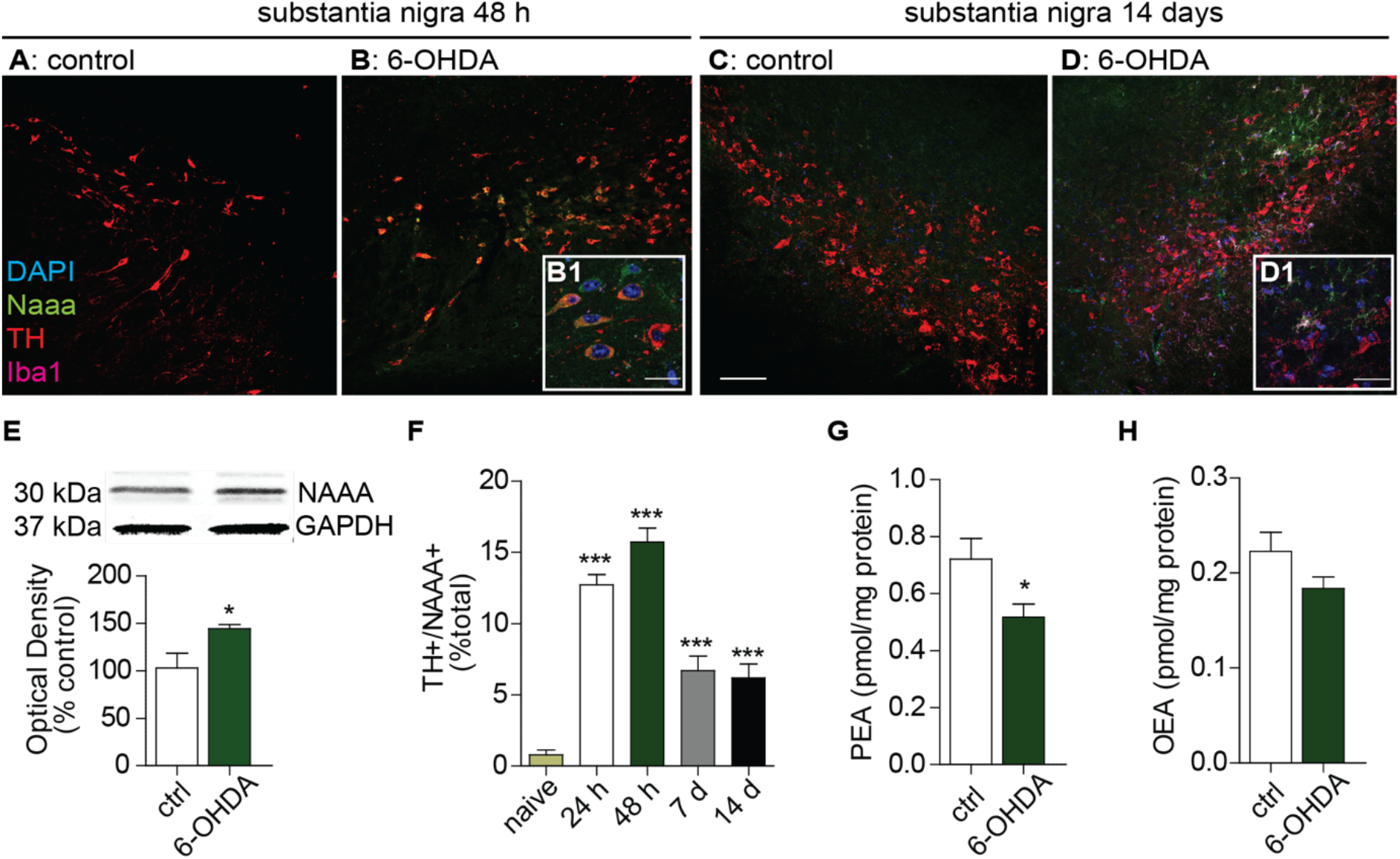
6-OHDA induces rapid and persistent NAAA expression in nigrostriatal dopamine neurons. • (A-D) Merged immunofluorescence images for NAAA (green), TH (red) and Iba-1 (magenta) in tissue sections of SN pars compacta: (A, C) control and (B, D) lesioned (ipsilateral and contralateral, respectively, to the 6-OHDA injection site); cell nuclei were stained with 4’,6-diamidino-2-phenylindole (DAPI, blue). Images were collected 48 h (A, B) or 14 days (C-D) after 6-OHDA injection. NAAA^+^/TH^+^ co-localization is shown in yellow, NAAA^+^/Iba-1^+^ co-localization is shown in white. Scale bar, 50 μm. (B1, D1) High-magnification images of NAAA^+^ cells that co-express (B1) TH or (d1) Iba-1. Scale bar, 10 μm. • (E) Western blot quantification of NAAA in midbrain fragments containing the SN: top, representative blot; bottom, densitometric quantification. Glyceraldehyde 3-phosphate dehydrogenase (GAPDH) was used for normalization. *P<0.05, two-tailed Student’s *t*-test (n=5). • (F) Number of neurons co-expressing TH and NAAA, shown as percent of total TH^+^ neurons in SN pars compacta. ***P<0.001, one-way ANOVA with Bonferroni post hoc test (n=4). • (G,H) Quantification of lipid content after 6-OHDA injection (G) PEA and (H) OEA in midbrain fragments containing the SN. *P<0.05, two-tailed Student’s *t*-test (n=5).

### Genetic or pharmacological NAAA ablation attenuates 6-OHDA-induced neurotoxicity

To investigate NAAA’s possible roles in 6-OHDA-induced neurotoxicity, we first used genetically modified mice that express the protein in a frame-shifted catalytically inactive form (*Naaa*^-/-^ mice). The mutants constitutively lack NAAA mRNA, protein, and enzyme activity (Supplementary Fig. 8), and do not compensate for this absence with altered transcription of isofunctional enzymes (fatty acid amide hydrolase) (Piomelli & Mabou Tagne, 2022), PEA-producing enzymes (*N*-acyl-phosphatidylethanolamine phospholipase D) (Okamoto *et al*, 2004) or other lysosomal lipid hydrolases (acid ceramidase and ß-glucocerebrosidase) (Park & Schuchman, 2006; Do *et al*, 2019) (Supplementary Fig. 8). As shown in Figure 5, homozygous NAAA deletion protected male mice from the cellular, neurochemical, and behavioral consequences of 6-OHDA injection. Three weeks after toxin administration, comparison with wild-type littermates showed that *Naaa^-/-^* mice had (*i*) improved survival of nigral TH^+^ neurons (Fig. 5A); (*ii*) higher striatal levels of dopamine (Fig. 5B) and dopamine metabolites (Supplementary Fig. 9); and (*iii*) greater density of striatal TH^+^ fibers (Fig. 5C). Moreover, *Naaa^-/-^* mice displayed *(iv)* prolonged latency to fall in the rotarod performance test (Fig. 5D); (*v*) attenuated motor responses (contralateral rotations) to the dopaminergic agonist apomorphine (0.1 mg/kg, subcutaneous; Fig. 5E); and (*vi*) lower mortality rate (Fig. 5F). Importantly, heterozygous *Naaa*^+/-^ mice were also resistant to 6-OHDA, but less so than their homozygous littermates (Fig. 5A-F). The gene dose-dependent neuroprotection caused by NAAA deletion points to a role for this enzyme in the control of dopamine neuron survival.

**Figure 4.**
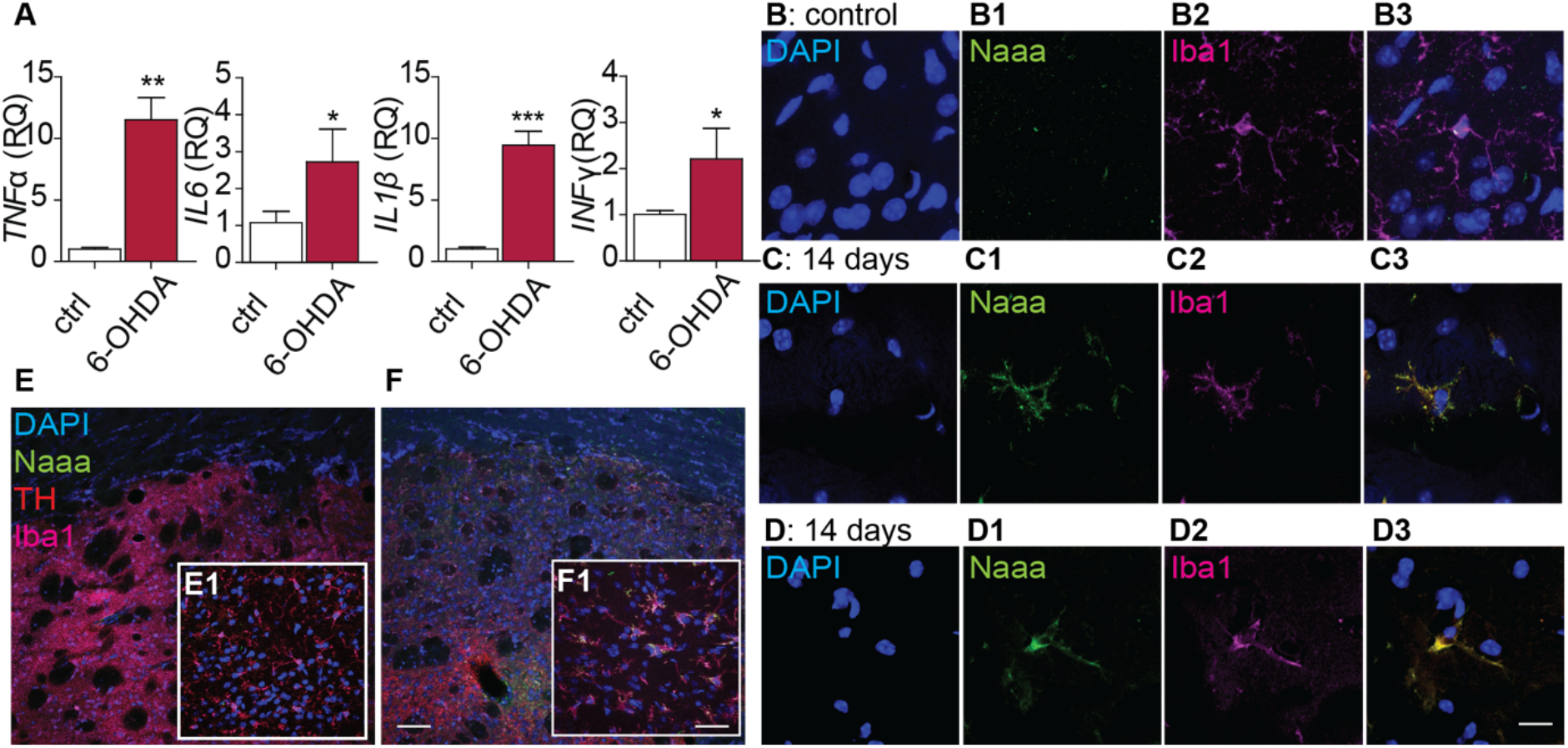
6-OHDA induces delayed NAAA expression in striatal microglia. • (A) Transcription of pro-inflammatory cytokines (tumor necrosis factor-a, TNF-a; interleukin 6, IL-6; interleukin-1b, IL-1b; and interferon-g, IFN-g) in control (open bars) and lesioned (red bars) striatum 14 days after 6-OHDA injection. ***P<0.001, **P<0.01, *P<0.05, twotailed Student’s *t*-test (n=5). • (B-D) Immunofluorescence images of sections from (B) intact and (C, D) lesioned striatum: cell nuclei are stained with DAPI (blue); NAAA is shown in green, Iba-1 in magenta; merged NAAA/Iba-1 signals are shown in yellow. Scale bar, 10 μm. • (E,F) Immunofluorescence images of sections from (E) intact and (F) lesioned striatum. Scale bar, 50 μm. • (E1, F1) High-magnification images of Iba-1^+^ microglial cells in sections from (E1) intact and (F1) lesioned striatum. Scale bar, 10 μm.

**Figure 5.**
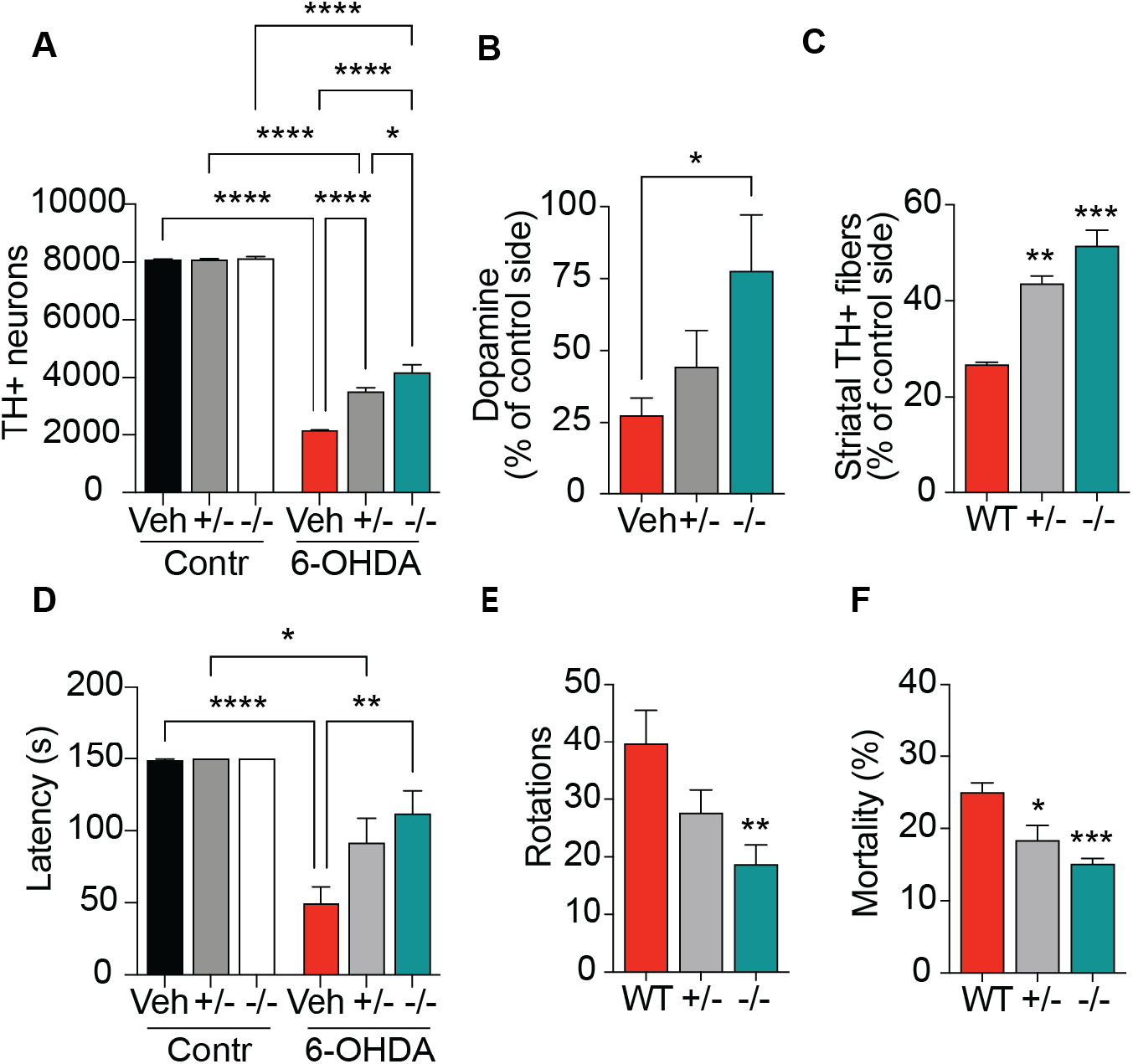
Genetic NAAA deletion protects mice from 6-OHDA-induced neurotoxicity. • (A) Number of TH^+^ neurons in SN pars compacta. ****P<0.0001, *P<0.01 two-way ANOVA, Bonferroni post hoc test (n=4). • (B) Striatal dopamine content. *P<0.05, compared to intact side one-way ANOVA, Tuckey post hoc test (n=4). • (C) Density of striatal TH^+^ fibers, expressed as percent of intact side. ***P<0.001, **P<0.01, one-way ANOVA and Tuckey post hoc test (n=6). • (D) Performance in the rotarod test (latency to fall, s); naïve mice (no 6-OHDA injection) are shown for comparison. ****P<0.0001, **P<0.01, *P<0.05, two-way ANOVA with Bonferroni post hoc test (n=8). • (E) Apomorphine-induced rotations. **P<0.01 compared to WT vehicle-treated mice, oneway ANOVA with Tuckey post hoc test (n=8). • (F) Mortality rate. ***P<0.001, *P<0.05, compared to WT mice, one-way ANOVA with Tuckey post hoc test (n=16). Measurements were performed 21 days after 6-OHDA injection.

Supporting this conclusion, we found that subchronic treatment with the systemically active NAAA inhibitor ARN19702 (Migliore *et al*, 2016) phenocopied genetic NAAA deletion. In male wild-type mice exposed to 6-OHDA, a 3-week twice-daily regimen with ARN19702 (30 mg/kg, intraperitoneal, starting the day of 6-OHDA administration) exerted a set of neuroprotective effects that included, in the SN, enhanced survival of TH^+^ neurons (Fig. 6A) and, in the dorsolateral striatum, increased dopamine and dopamine metabolite content (Fig. 6B, Supplementary Fig. 10) and density of TH^+^ fibers (Fig. 6C). Additionally, the NAAA inhibitor prolonged the latency to fall in the rotarod test (Fig. 6D), attenuated motor responses to apomorphine (Fig. 6E) and decreased animal mortality (Fig. 6F). Thus, as seen with *Naaa* deletion, pharmacological NAAA inhibition enhances dopamine neuron survival and ameliorates parkinsonian symptoms in mice treated with 6-OHDA.

**Figure 6.**
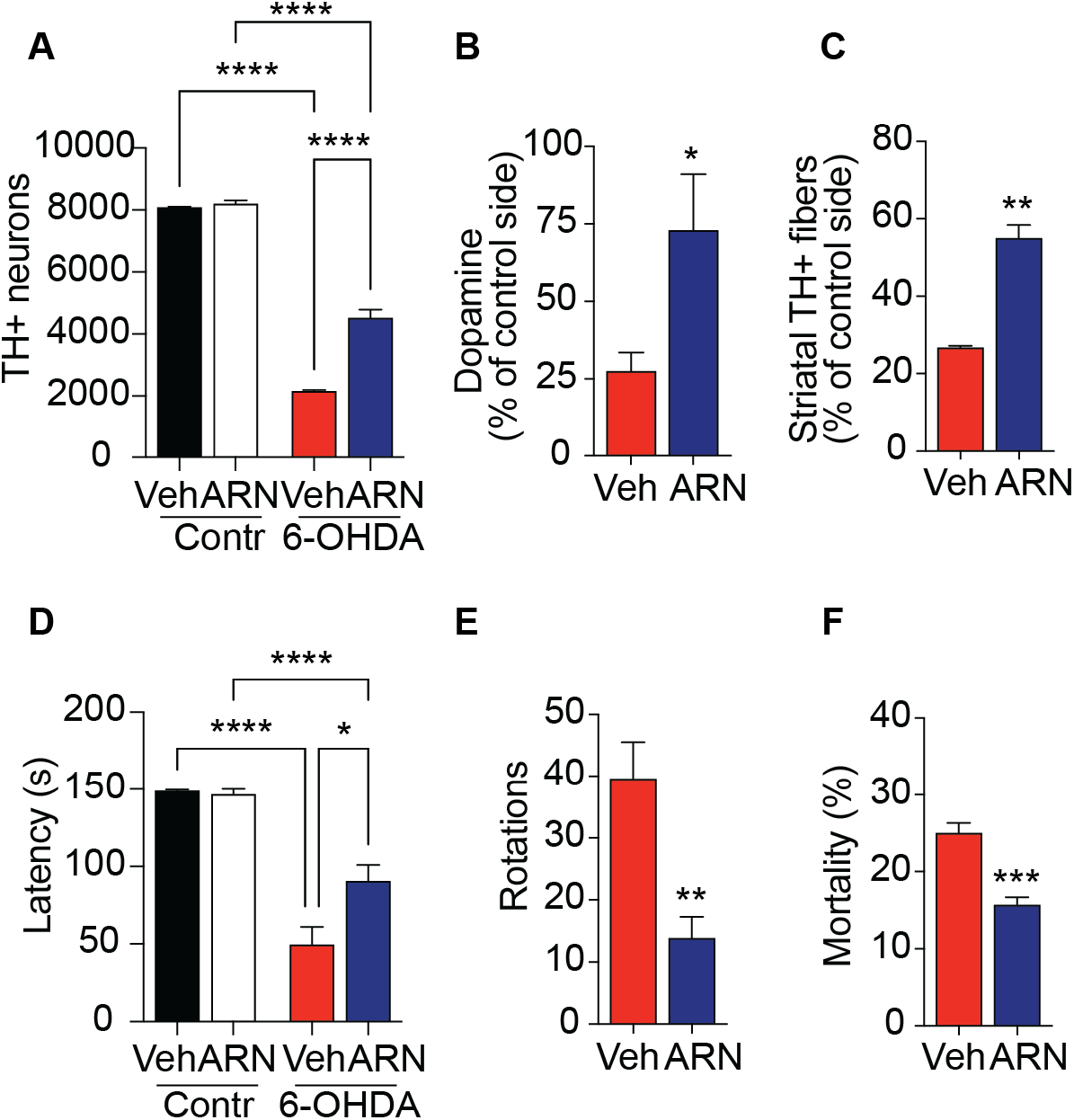
Pharmacological NAAA inhibition protects mice from 6-OHDA-induced neurotoxicity. • (A) Number of TH^+^ neurons in SN pars compacta. ****P<0.0001, two-way ANOVA, Bonferroni post hoc test (n=4). • (B) Striatal dopamine content. *P<0.05, compared to intact side Student’s *t* test (n=4). • (C) Density of striatal TH^+^ fibers, expressed as percent of intact side. **P<0.01, Student’s *t* test (n=6). • (D) Performance in the rotarod test (latency to fall, s); naïve mice (no 6-OHDA injection) are shown for comparison. ****P<0.0001, *P<0.05, two-way ANOVA with Bonferroni post hoc test (n=8). • (E) Apomorphine-induced rotations. **P<0.01, Student’s *t* test (n=8). • (F) Mortality rate. ***P<0.001, Student’s *t* test (n=16). Measurements were performed 21 days after 6-OHDA injection.

### Genetic or pharmacological NAAA ablation attenuates MPTP-induced neurotoxicity

Next, we asked whether the neuroprotective effects of NAAA removal might generalize to MPTP-induced neurotoxicity (Jackson-Lewis & Przedborski, 2007). Within 7 days of administration, MPTP produced in female wild-type mice a substantial loss of SN dopamine neurons (Fig. 7A) and a decrease in striatal dopamine and dopamine metabolite levels (Fig. 7B and Supplementary Fig. 11) and TH+ fiber density (Fig. 7C). These morphological and neurochemical events were accompanied by a worsened performance in the rotarod test (Fig. 7D) and marked animal mortality (Fig. 7E). Importantly, the effects of MPTP were substantially attenuated in *Naaa^-/-^* mice (Fig. 7A-E). As seen in the 6-OHDA model, treatment with ARN19702 (30 mg/kg, twice daily, intraperitoneal, starting the day before MPTP administration) phenocopied *Naaa* deletion, improving dopamine neuron survival (Fig. 8A), increasing striatal dopamine content (Fig. 8B) and fibers (Fig. 8C), normalizing behavior in the rotarod test (Fig. 8D) and curbing animal mortality (Fig. 8E). Collectively, the results suggest that NAAA-regulated PEA signaling protects nigral dopamine neurons against 6-OHDA- and MPTP-induced toxicity.

**Figure 7.**
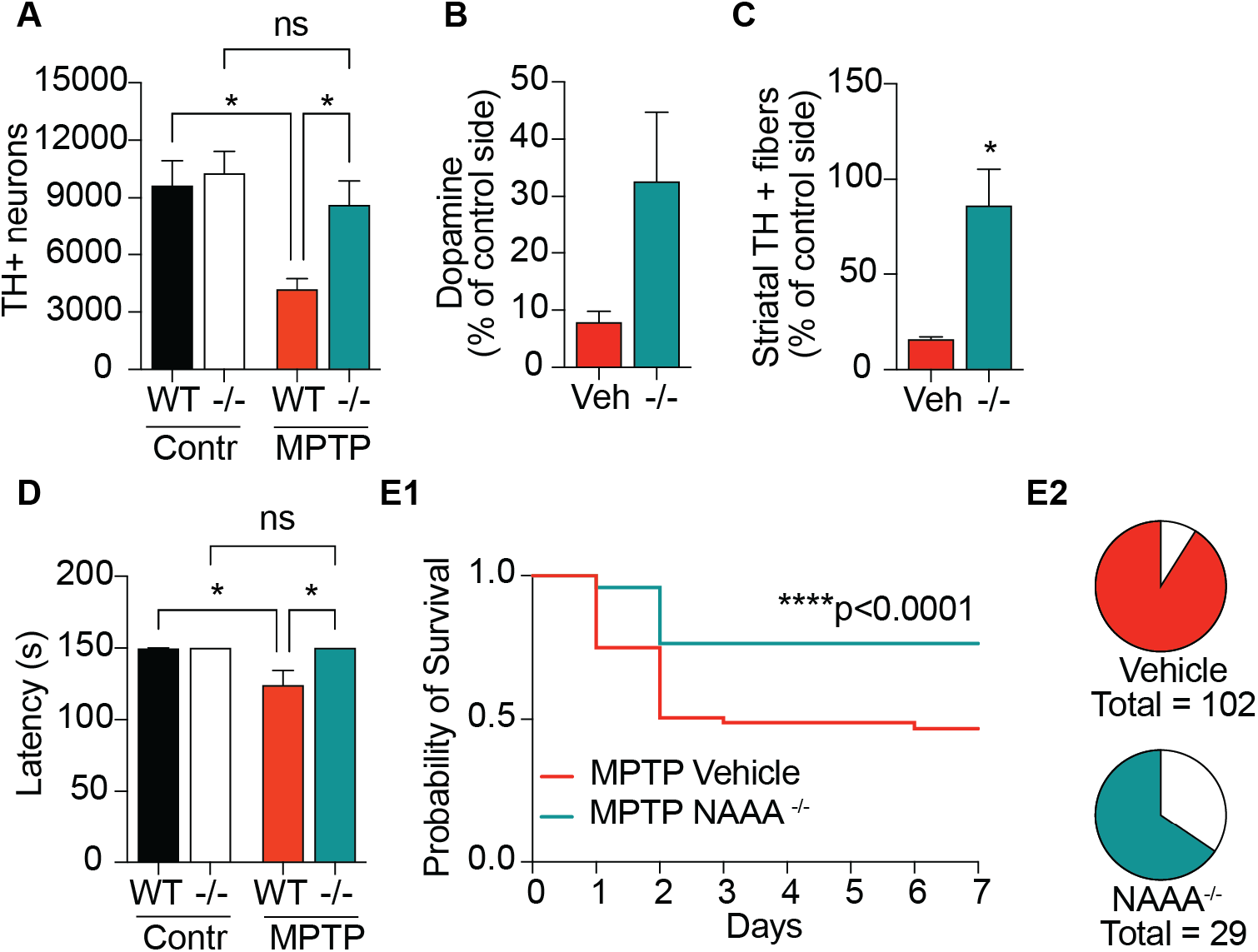
Genetic NAAA deletion protects mice from MPTP-induced neurotoxicity. • (A) Number of TH^+^ neurons in SN pars compacta. two-way ANOVA, Bonferroni post hoc test (n=3-5). • (B) Striatal dopamine content (n=3-4). • (C) Density of striatal TH^+^ fibers, expressed as percent of controls. *P<0.05, Student’s *t* test (n=3-4). • (D) Performance in the rotarod test (latency to fall, s); naïve mice (no MPTP injection) are shown for comparison. *P<0.05, two-way ANOVA with Bonferroni post hoc test (n=3-10). • (E1) Survival rate. ****P<0.001, Log-rank test (n=29-102). • (E2) Total number of animals survived after MPTP administration. Measurements were performed 7 days after MPTP injection.

**Figure 8.**
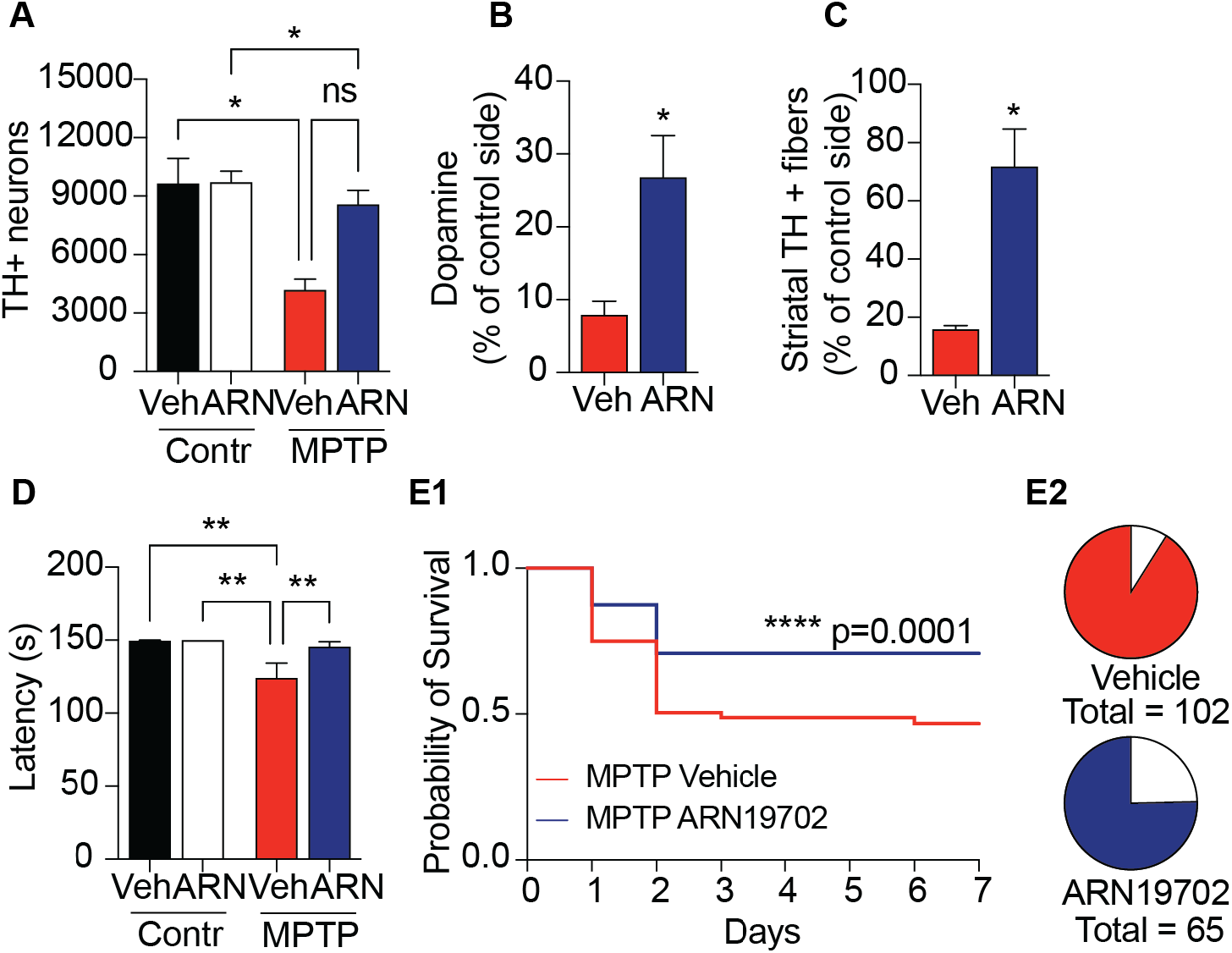
Pharmacological NAAA inhibition protects mice from MPTP-induced neurotoxicity. • (A) Number of TH^+^ neurons in SN pars compacta. two-way ANOVA, Bonferroni post hoc test (n=3-5). • (B) Striatal dopamine content. *P<0.05, Student’s *t*-test (n=3-4). • (C) Density of striatal TH^+^ fibers, expressed as percent of controls. *P<0.05, Student’s *t* test (n=3-5). • (D) Performance in the rotarod test (latency to fall, s); naïve mice (no MPTP injection) are shown for comparison. **P<0.01, two-way ANOVA with Bonferroni post hoc test (n=8-15). • (E1) Survival rate. ****P<0.001, Log-rank test (n=65-102). • (E2) Total number of animals survived after MPTP administration. Measurements were performed 7 days after MPTP injection.

### Mechanistic insights

To gain insights into possible mechanisms through which NAAA might influence neuronal survival, we compared the nigral proteomes of *Naaa*^-/-^ and wild-type mice 48 h after 6-OHDA administration, when NAAA expression is maximal in dopamine neurons but still undetectable in microglia. Analyses of lesioned SN tissue identified 607 differentially expressed proteins, most of which (603) were overrepresented in *Naaa*^-/-^ mice compared to wild-type controls (Supplementary Table 9). Annotation using the Database for Annotation, Visualization, and Integrated Discovery (DAVID) (Huang *et al*, 2009) showed that proteins upregulated in *Naaa^-/-^* mice included many components of oxidative phosphorylation (Fig. 9A), carbon metabolism (Fig. 9C) and lipid and glycan metabolism (Fig. 9D). Similarly, Gene Ontology (GO) analysis revealed a substantial enrichment in proteins that are (*i*) part of the Krebs’ cycle, oxidative phosphorylation, and glycolysis (GO category ‘biological process’); *(ii)* involved in ATP synthesis, proton transport and redox function (‘molecular function); and *(iii)* localized to the mitochondrial inner membrane and cytoskeleton (‘cellular component’) (Supplementary Fig. 15). In contrast with the profound changes observed in lesioned SN, parallel studies of intact SN found only minor differences between *Naaa*^-/-^ and wild-type mice (Fig. 9B, E, F). The marked impact of NAAA deletion on cellular bioenergetics pathways led us to hypothesize that members of the PGC-1 family, which are crucial upstream regulators of those pathways and have been implicated in dopamine neuron survival and the pathogenesis of PD (Jiang *et al*, 2016; Zheng *et al*, 2010; Su *et al*, 2015; Piccinin *et al*, 2021), might be involved. Supporting this conclusion, a closer inspection of the proteomics data identified type-ß peroxisome proliferator-activated receptor-γ coactivator-1 (PGC-1ß) as one of the proteins deregulated by 6-OHDA (Fig. 10A, Supplementary Table 9: *replicate 2, line 641*). Western blot analyses confirmed that PGC-1ß is overrepresented in lesioned SN of *Naaa*^-/-^ mice, relative to lesioned SN of wild-type controls (Fig. 10 B, C), and further showed that another key member of the PGC-1 family, PGC-1 α, is also subject to NAAA regulation (Fig. 10 D, E). It is noteworthy that PGC-1α and ß were underrepresented in lesioned SN of wild-type mice – as seen in persons with PD (Zheng *et al*, 2010) – whereas, in *Naaa*^-/-^ mutants, they were both constitutively elevated and did not change in response to 6-OHDA (Fig. 10 B-E).

**Figure 9.**
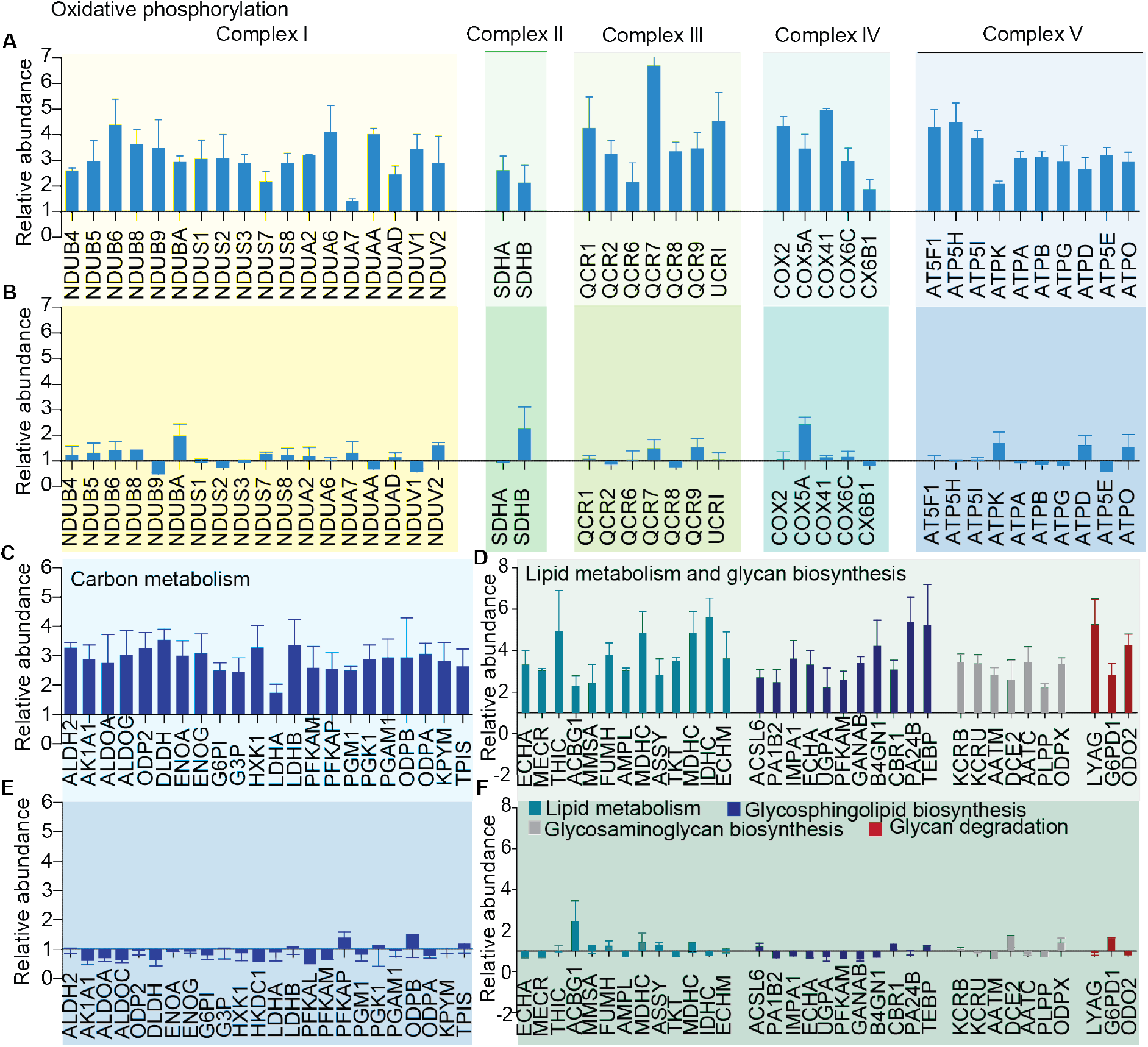
*Naaa^-/-^* mice differentially regulate bioenergetics pathways in response to 6-OHDA. • (A,C,E) Comparative proteomics analysis of lesioned SN *(Naaa^-/-^* versus wild-type, WT): (A) oxidative phosphorylation; (C) carbon metabolism; and (E) lipid and glycan metabolism. • (B,D,F) Comparative proteomics analysis of intact SN *(Naaa^-/-^* versus WT): (B) oxidative phosphorylation; (D) carbon metabolism; and (F) lipid and glycan metabolism.

**Figure 10.**
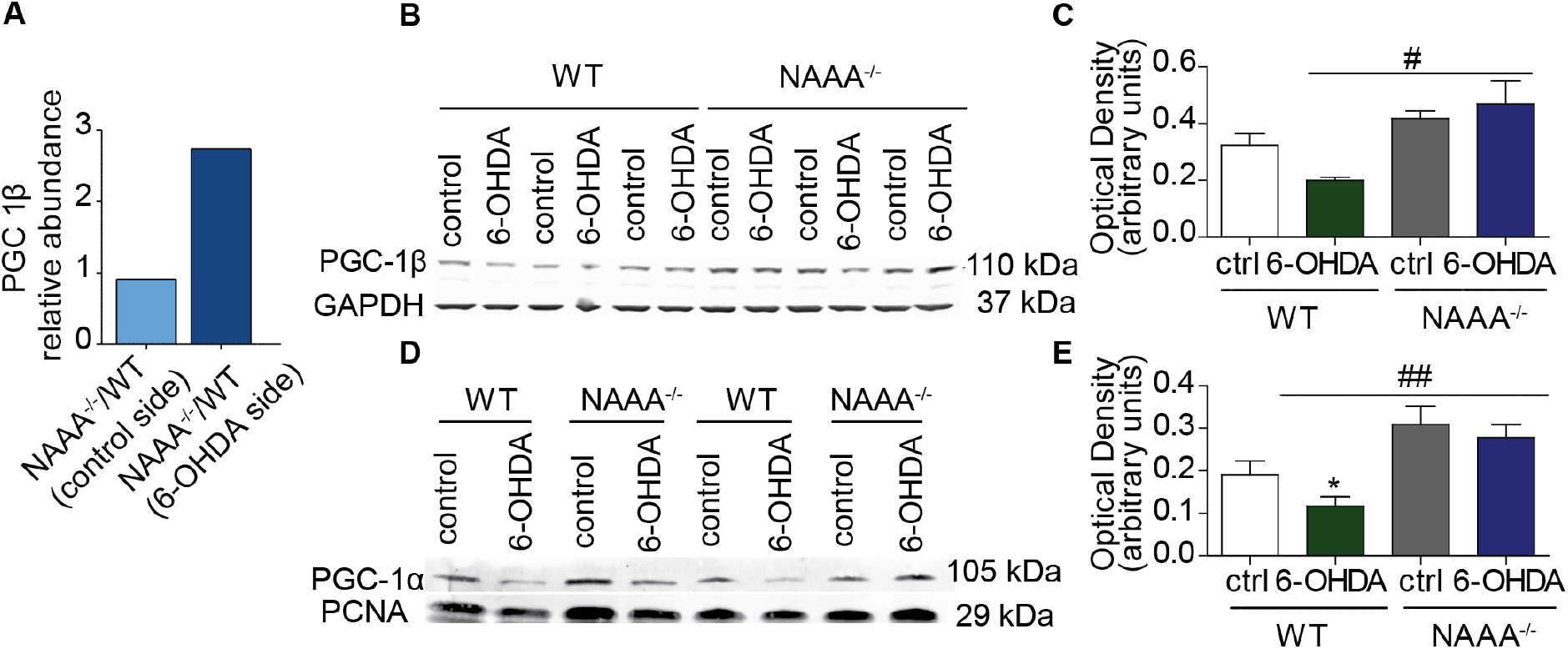
6-OHDA suppresses expression of bioenergetics pathways in a NAAA-dependent manner. • (A) Relative abundance *(Naaa^-/-^* versus WT) of PGC-1β in control (light blue) and lesioned (dark blue) SN. • (B-E) Representative western blots and densitometric quantification of (C) PGC1-ß in nigral homogenates, and (E) PGC1-α in nuclear fractions of SN tissue from WT and *Naaa^-/-^* mice. GAPDH and proliferating cell nuclear antigen (PCNA) were used for normalization. *P<0.05 compared to wild-type (WT) contralateral side; #P<0.05 and ## P<0.01, compared to lesioned side of WT/veh mice, two-way ANOVA, Bonferroni post hoc test (n=4).

## DISCUSSION

The study’s objective was to determine whether NAAA, a lysosomal cysteine hydrolase previously implicated in inflammation and nociception (Ueda *et al*, 2010; Piomelli *et al*, 2020) might contribute to the pathogenesis of PD. We carried out three experiments to test this idea. First, we measured NAAA mRNA and protein levels in blood-derived exosomes and frozen brain specimens of persons with PD. Consistent with an earlier gene-array survey (Papapetropoulos *et al*, 2006), the analyses found evidence for elevated premortem and postmortem NAAA expression in these patients, compared to age- and sex-matched controls. This finding prompted us to ask whether a link might exist between NAAA-regulated PEA signaling and dopamine neurons damage – a neuropathological hallmark of PD (Vijiaratnam *et al*, 2021). Thus, in a second set of experiments we investigated the effects of 6-OHDA and MPTP, two neurotoxins that cause PD-like pathology in rodents and humans (Dauer & Przedborski, 2003; Langston *et al*, 1983), on NAAA expression. Incubation of human SH-SY5Y cells with either toxin produced a substantial increase in NAAA mRNA and protein along with a decline in cellular PEA content. Similar changes were observed in mouse midbrain, where intrastriatal injections of 6-OHDA stimulated NAAA expression in both nigrostriatal dopamine neurons and microglia recruited to the sites of injury. These observations motivated one last set of experiments, in which we examined whether removal of the *Naaa* gene or inhibition of NAAA activity might attenuate dopamine neuron death caused in mice by administration of 6-OHDA or MPTP. We found that both genetic and pharmacological NAAA ablation enhanced dopamine neuron viability and countered PD-like symptoms elicited by the neurotoxins. Collectively, the results identify NAAA as a molecular checkpoint for the survival of nigrostriatal dopamine neurons and a potential target for PD modification.

It is possible that NAAA-regulated PEA signaling influences dopamine neuron viability by modulating microglia activity and neuroinflammation (Migliore *et al*, 2016; Pontis *et al*, 2020; Sgroi *et al*, 2021). Our findings point, however, to a more complex scenario. In mice treated with 6-OHDA, the rise in NAAA expression is initially confined to dopamine neurons (Fig. 3) and only later extends to reactive microglia (Fig. 4). The ‘neuronal’ phase of NAAA induction begins within 24 h of exposure to the toxin (Fig. 3 and Supplementary Fig. 4) – that is, well before the appearance of overt signs of neuroinflammation – and is part of a broad program that includes the suppression of protein networks responsible for mitochondrial biogenesis and cellular respiration (Supplementary Fig. 5 and 6). This suggests that PEA-mediated signaling at PPAR-α may protect dopamine neurons, at least in part, by preserving their bioenergetic potential – as recently proposed for spinal cord neurons after peripheral organ injury (Fotio *et al*, 2021b) – and that pathological NAAA induction might undermine this protection. Supporting this view, our results also indicate that NAAA deletion *(i)* prevents the disruptive effects of 6-OHDA on bioenergetics pathways (Fig. 9); and *(ii)* normalizes expression of the transcription coactivators, PGC-1α and PGC-1ß, which are suppressed after 6-OHDA challenge (Fig. 10 C). PGC-1α and PGC-1ß serve pivotal functions in metabolic control (Handschin & Spiegelman 2006) and, importantly, have been implicated in dopamine neuron survival (Jiang *et al*, 2016) and the pathogenesis of PD (Zheng *et al*, 2010; Su *et al*, 2015; Piccinin *et al*, 2021; Das et al, 2021; Jamwal *et al*, 2021). Similar roles have been ascribed to PPAR-α (Gottschalk *et al*, 2021; Avagliano *et al*, 2016), PEA’s primary receptor and obligatory PGC-1α partner (Handschin & Spiegelman 2006).

The relevance of our results to sporadic PD in humans requires consideration. We present evidence that NAAA expression is elevated in postmortem brain cortex from persons with advanced PD (average age of death: 76.5 ± 4.9 years; disease duration: 15.3 ± 8.3 years; mean ± SD) as well as in blood-derived exosomes from persons at stages I and II of the disease, according to Hoehn and Yahr (average age at collection: 54.7 ± 14.8 years; disease duration: 3.58 ± 2.6 years). This finding is consistent with a role for NAAA in the pathogenesis of PD, but also raises two critical questions: what genetic or environmental stimuli might be responsible for triggering NAAA elevation? And, when could such elevation occur in an individual’s lifespan and in which organ system(s)? The heterogeneous etiology, long prodromic phase, and slow progression of PD (Bloem *et al*, 2021) make answering these questions especially arduous. One possible scenario, however, is that head injury or exposure to mitotoxic pollutants, two established risk factors for PD (Bloem *et al*, 2021), may enhance directly or indirectly NAAA expression, as suggested by the enzyme’s inducible nature (Fotio *et al*, 2021b). Alternatively, or possibly complementarily, NAAA upregulation may result from neuronal damage caused by accumulation of insoluble α-synuclein aggregates, a neuropathological hallmark of PD (Vijiaratnam *et al*, 2021). Given the multisystem spread of PD pathology (Chahine *et al*, 2020), both events might take place in brain, peripheral organs, or both.

The study has two main weaknesses. First, human tissue analyses included a relatively small group of subjects (PD: 54; controls: 58) which was composed primarily by women (PD: 39; controls: 45). It will be important to address both sample size limitation and sex bias in a larger cohort, not only to replicate (or not) the initial findings but also to explore the possible value of NAAA in circulating exosomes as a peripheral biomarker of PD progression. Second, our animal studies relied on two toxin models which, though well-characterized and widely used, do not recapitulate the slow progressive nature of sporadic PD. A possible alternative, which will be evaluated in future work, is offered by the α-synuclein pre-formed fibril model, which is gaining traction after its recent standardization (Polinski *et al*, 2018). Despite these limits, the present report provides multiple lines of evidence indicating that NAAA plays a critical role for in the control of nigrostriatal dopamine neuron survival and might thus offer a druggable target for therapeutic intervention in PD.

## MATERIALS AND METHODS

### Ethics statement

Investigations were conducted in accordance with the Declaration of Helsinki as well as with international guidelines. Moreover, they were approved by each of the authors’ institutional review boards.

### Chemicals

6-OHDA hydrochloride, MPP^+^ iodide, chloral hydrate, ketamine, xylazine, paraformaldehyde, MPTP, dopamine, serotonin, homovanillic acid (HVA), 3,4-dihydroxyphenylacetic acid (DOPAC), ascorbic acid, and [^2^H_5_]-benzoyl chloride were purchased from Sigma Aldrich (Saint Louis, MO). [^2^H_4_]-PEA and [^2^H_4_]-OEA were from Cayman Chemical (Ann Arbor, MI, USA). Tandem Mass Tag™ 6-plex (TMTsixplex™) reagents kits for isotopic labeling were from Thermo Fisher Scientific (Waltham, MA, USA). All analytical solvents were of the highest grade, and were obtained from Honeywell (Muskegon, MI) or Sigma-Aldrich. ARN19702 was synthetized as described (Migliore *et al*, 2016).

### Cell cultures

SH-SY5Y cells were purchased from Sigma Aldrich and cultured at 37°C and 5% CO_2_ in Dulbecco’s Modified Eagle’s Medium (DMEM) (Euroclone, Milan, Italy) supplemented with 10% fetal bovine serum (FBS, Thermo Fisher Scientific), L-glutamine (2 mM) and antibiotics (Euroclone). Cells were incubated with 6-OHDA (100 μM), MPP^+^ (2 mM) or their respective vehicles (6-OHDA: saline containing 0.2% ascorbic acid; MPP^+^: complete DMEM) for the indicated times.

### Postmortem human brain specimens

Postmortem brain cortex samples were obtained from the Banner Sun Health Research Institute (Hoss *et al*, 2016; Beach *et al*, 2015). Specimens from 46 control subjects and 43 patients with postmortem interval 1.25 – 4.83 h were used to prepare mRNA and protein extracts and to conduct morphological analyses, as described below. Detailed demographic and clinical information are provided in Supplementary Table 1.

### Human study subjects

Potential study participants were assessed at the Laboratory of Neuropsychiatry of the I.R.C.C.S. Santa Lucia Foundation in Rome. Inclusion criteria for PD patients were: (i) diagnosis of idiopathic PD according to the international guidelines (Folstein *et al*, 1975) (ii) Mini-Mental State Examination (MMSE) score ≥26 and no dementia according to the Movement Disorder Society (MDS) clinical diagnostic criteria (Emre *et al*, 2007). Exclusion criteria were (i) presence of major non stabilized medical illnesses (i.e., non-stabilized diabetes, obstructive pulmonary disease or asthma, hematologic/oncologic disorders, vitamin B12 or folate deficiency, pernicious anemia, clinically significant and unstable active gastrointestinal, renal, hepatic, endocrine or cardiovascular disorders and recently treated hypothyroidism; (ii) known or suspected history of alcoholism, psychoactive drug dependence, head trauma, and mental disorders (apart from mood or anxiety disorders) according to the DSM-IV criteria (Bell, 1994); (iii) history of neurological diseases other than idiopathic PD; (iv) unclear history of chronic dopaminergic treatment responsiveness; and (v) MRI scans lacking signs of focal lesions as computed according to a semi-automated method (McKhann *et al*, 2011) [minimal diffuse changes or minimal lacunar lesions of white matter (WM) were, however, allowed]. All PD patients included in the study were under stable dopaminergic therapy and were at stage I or II of the disease. Furthermore, they were not receiving Deep Brain Stimulation and were not under continuous dopaminergic stimulation by subcutaneous apomorphine or intrajejunal L-DOPA. Inclusion criteria for control subjects were: (i) vision and hearing sufficient for compliance with testing procedures; (ii) laboratory values within the appropriate normal reference intervals; and (iii) neuropsychological domain scores above the cutoff scores, corrected for age and educational level, identifying normal cognitive level in the Italian population. Exclusion criteria were: (i) dementia diagnosis, according with DSM-V criteria or MCI according with Petersen criteria and confirmed by a comprehensive neuropsychological battery; and (ii) MMSE score <26 according with standardized norms for the Italian population (Dellasega & Morris, 1993). After blood sampling, the patients’ identities were masked by a random code and all personal information was deleted. All the participants or related caretakers gave their written consent to the enrollment. The study was approved by the Ethical Committee of I.R.C.C.S. Santa Lucia Foundation (CE-AG4-Prog.149) and by the Italian Ministry of Health, which also provided financial support (Grant RF-2013-02359074). Sociodemographic and clinical characteristics of study participants are reported in Supplementary Table 2.

### Human blood collection

Blood samples (10 mL) were collected after overnight fasting (11 ± 1 h) from forearm veins into BD Vacutainer tubes containing EDTA (Beckton Dickinson, Franklin Lakes, NJ). Plasma was prepared by centrifugation at 4°C and was stored at −80°C until analyses.

### Animals

All procedures were performed in accordance with the Ethical Guidelines of the European Union (directive 2010/63/EU of 22 September 2010) and the Italian Ministry of Health or were approved by the Institutional Animal Care and Use Committee of the University of California, Irvine. The experiments were carried out in strict accordance with the National Institutes of Health guidelines for care and use of experimental animals. C57BL/6 mice (20-35 g) were purchased from Charles River (Wilmington, MA, USA). Male mice were used for 6-OHDA experiments, female mice were used for MPTP studies. B6N-Atm1BrdNaaatm1a(KOMP)wtsi/WtsiH mice (*Naaa^-/-^*) were generated at the Welcome Trust Sanger Institute and obtained from Jackson Labs (Bar Harbor, ME, USA). *Naaa^-/-^* mice are viable and fertile, and do not display overt behavioral abnormalities. Genotyping was performed by polymerase chain reaction (PCR) using DNA (30 ng) extracted from tail clips taken at 21 days of age. PCR reactions were run using PCR SUPERMIX (Invitrogen, Carlsbad, CA, USA) as shown in Supplementary Table 15. All mice were group-housed in ventilated cages and had free access to food and water. They were maintained under a 12h light/dark cycle (lights on at 8:00 am) at controlled temperature (21 ± 1°C) and relative humidity (55% ± 10%). All efforts were made to minimize animal suffering and to use the minimal number of animals required to produce reliable results.

### 6-OHDA and MPTP-induced neurotoxicity

Mice were anesthetized with a mixture of ketamine (87.5 mg/kg) and xylazine (12.5 mg/kg; 0.1 mL per 20 g body weight, intraperitoneal, i.p.) and placed in a stereotaxic frame with a mouse adaptor (Kopf Instruments, Tujunga, CA, USA). 6-OHDA was dissolved at a fixed concentration of free base (3.2 mg/mL) in ice-cold saline containing ascorbic acid (0.02%). Two 6-OHDA injections (1 μL each) were made using a 33-gauge Hamilton syringe needle (Hamilton, Reno, NV, USA) at the following brain coordinates: (*i*) AP=+1.0, L=-2.1, DV=-2.9; and (*ii*) AP=+0.3, L=-2.3, DV=-2.9 (Watson & Paxinos, 2012). Injections were performed at a rate of 0.5 μL/min and 2 min were allowed for the toxin to diffuse. MPTP was dissolved at the concentration of 18 mg/kg in ice cold saline and administered by i.p. injection every 2h for a total of 4 doses over an 8-h period in a single day, as previously described (Jackson-Lewis & Przedborski, 2007).

### Drug treatments

ARN19702 was dissolved in a vehicle composed of sterile saline containing PEG-400 (15%, vol/vol) and Tween (15%). The drug (30 mg/kg, i.p.) or its vehicle was injected twice a day at approximately 8:00 a.m. and 6:00 p.m. Treatment started either on the day of 6-OHDA injection and was continued for the following 3 weeks, or the day before MPTP administration and lasted 7 days.

### Mouse tissue processing and immunohistochemistry

Mice were anaesthetized with chloral hydrate (450 mg/kg, i.p.) and perfused transcardially with ice-cold sterile saline (20 mL), followed by ice-cold paraformaldehyde [PFA, 4% in phosphate-buffered saline (PBS), 60 mL]. The brains were excised and stored in sucrose (25% in PBS) at 4°C. Three series of sections (thickness 40 μm) were collected in the coronal plane using a cryostat and stored at −20°C. Double and triple immunostaining protocols were performed by sequential incubation with primary antibodies (1:200; Abcam, Cambridge, UK) followed by secondary Alexa Fluor 488 antibodies (1:1000; Invitrogen). Images were collected using a Nikon A1 confocal microscope with a 60 x 1.4 numerical aperture objective lens. Quantification of colocalization was performed in double immunostained sections. A minimum of 50 cells (3 coverslips/mouse) was analyzed combining Alexa Fluor-488 conjugated and Alexa Fluor-633 conjugate secondary antibodies. Costes automatic threshold of green and magenta channels was selected. The background was subtracted from the original images, the actual images were analyzed. The assessment of colocalization of NAAA immunofluorescence (green channel) and tyrosine hydroxylase (TH) or Iba1 (magenta channel) immunofluorescence was performed using the NIH ImageJ Just Another Colocalization plug-in (JACoP) (available at http://rsb.info.nih.gov/ij/plugins/track/jacop.html). JACoP provides the Pearson R correlation coefficient (PC), Mander’s M1 and M2 coefficients and Li’s intensity correlation quotient (ICQ) coefficients for a pair of images. Graph Prism software was used to generate all the bar graphs and statistical analysis of the data. Results are presented as mean ± SEM of six mice.

Quantification of the percentage of TH^+^ neurons expressing NAAA was conducted on 6 sections per mouse, 4 mice per time point were analyzed. To determine whether a cell was positive to both the selected neurochemical markers, confocal images from of the same focal plane were acquired in the two channels and co-localization analysis was run as described above. The percentage of NAAA^+^ and TH^+^ cells over the total number of TH^+^ cells was quantified for each section and an average value was calculated for each mouse.

### Human tissue processing and immunohistochemistry

Postmortem human cortex samples were sectioned (thickness 40 μm) using a cryostat and stored at −20°C. Slices were fixed in icecold PFA (4% in PBS) and double immunostaining protocols were performed by sequential incubation with primary antibodies (Supplementary Table 16) followed by secondary Alexa Fluor antibodies (1:1000; Invitrogen). Images were collected using an Olympus (Tokyo, Japan) FV3000 confocal microscope with a 40 x 1.25 numerical aperture objective lens.

### Stereological measurements

Dopamine neurons of the SN were identified after TH staining followed by Alexa fluor 550/488 secondary antibody of midbrain regions with a 4x objective. The SN was outlined using the Paxinos and Franklin’s mouse brain atlas as reference. Quantitative estimation of TH^+^ neurons was performed in every 6th section. Briefly, unbiased sampling and blinded stereological counting were performed using the optical fractionator probe of the Stereo Investigator software (MBF Bioscience, Williston, VT, USA). Parameters used included a 60x oil objective, a counting frame size of 60 × 60, a sampling site of 100 × 100, a dissector height of 15 μm, 2 μm guard zones. The Gunder’s coefficient of error was less than 0.1. A total of 5 animals per group were used and 5 to 8 sections per animal were counted in the red/green channel.

### Exosome isolation

1 mL of human plasma was incubated with 0.5 mL of Invitrogen total exosome isolation reagent (Thermo Fisher Scientific) overnight at 4°C and the day after was centrifuged at 10,000 *x g* for 1h at 4°C. Pellets were resuspended in 2 volumes of radioimmunoprecipitation assay buffer (RIPA, 150 μL, 50 mM Tris-HCI pH 7.4, 1% NP40, 0.5% Na-deoxycholate, 0.1% SDS, 150 mM NaCI, 2mM EDTA), homogenized and sonicated. Protein concentration was measured using the bicinchoninic acid (BCA) method, according to manufacturer’s instructions (Thermo Fisher Scientific) and 30 μg of protein was used for western blot analyses. Cell media from 6-OHDA- or vehicle-treated SH-SY5Y cells were collected after 8 h of incubation and centrifuged at 800 *x g* for 10 min at 4°C. Supernatants were filtered using a 0.22 μm filter and centrifuged at 100,000 *x g* for 90 min at 4°C. Pellets were resuspended in 0.5 mL PBS containing a proteinase inhibitor cocktail (Thermo Fisher Scientific) and centrifuged again at 100,000 *x g* for 90 min at 4°C. Supernatants were collected and protein concentration measured using the BCA method. Thirty μg of protein were used for western blot analyses.

### Western blot analyses

Human cortex or mouse midbrain fragments containing the left or right SN were dissected under microscope guidance and homogenized in RIPA buffer (150 μL). Protein concentration was measured using BCA. PGC1-α was identified and quantified in nuclear-enriched fractions. Briefly, brain tissue dissected from control and lesioned hemispheres was homogenized in buffer [v/w=5:1; 10 mM Hepes-KOH (pH 7.4), 0.25 M sucrose, proteinase inhibitor cocktail (Sigma)]. Homogenates were centrifuged at 900 x *g* for 10 min. Pellets were collected, homogenized again, and centrifuged at 900 x *g* for 10 min, to yield the nuclear fraction (pellet). Proteins (30 μg) were denatured in sodium dodecyl sulfate (SDS, 8%) and β-mercaptoethanol (5%) at 95°C for 5 min. After separation by SDS-PAGE on a 4-15% gel under denaturing conditions, the proteins were electrotransferred to nitrocellulose membranes. The membranes were blocked with goat serum (5% in PBS) and incubated overnight with primary antibodies in I-block solution (Thermo Fisher Scientific), followed by incubation with the corresponding secondary antibody (Invitrogen) in tris-buffered saline at room temperature for 1h. Fluorescence gel scanning was performed with FLA-9000 Starion (Fujifilm Life Science, USA), or Fluorchem R (ProteinSimple, San Jose, CA, USA). Antibody sources and dilutions are shown in Supplementary Table 16.

### Lipid extraction and liquid chromatography-mass spectrometry (LC/MS) analysis

Cell and tissue levels of PEA and OEA were measured as described (Zhu *et al*, 2011). Briefly, snap-frozen cell pellets or pre-weighed tissues were homogenized in methanol (1 mL) containing [^2^H_4_]-PEA and [^2^H_4_]-OEA (Cayman Chemical) as internal standards. Lipids were extracted with chloroform (2 vol), and organic phases were washed with water (1 vol), collected, and dried under nitrogen. The organic extracts from cells were reconstituted in methanol (0.1 mL) and fractionated by silica gel column chromatography (Astarita *et al*, 2009). PEA and OEA were eluted with chloroform/methanol (9:1, v/v). The eluates were evaporated under nitrogen and reconstituted in 75 μL of methanol/chloroform (9:1, v/v). LC/MS analyses were conducted on a Xevo TQ LC-MS/MS system equipped with a BEH C18 column, using a linear gradient of acetonitrile in water. Quantification was performed monitoring the following MRM transitions (parent *m/z* > daughter *m/z,* collision energy eV): OEA 326>62,20; [^2^H_4_]-OEA 330>66,20; PEA 300>62,20; [^2^H_4_]-PEA 304>66,20. Analyte peak areas were compared to a standard calibration curve (1 nM to 10 μM).

### NAAA activity assay

Lysosomal-enriched samples (50 mg of enriched samples obtained as previously described (Bonezzi *et al*, 2016)) were incubated in assay buffer (pH 4.5; 150 mM NaCl, 100 mM trisodium citrate dihydrate, 100 mM sodium phosphate monobasic, NaH_2_PO_4_, 0.1% Nonidet P-40, and 3 mM dithiothreitol) in a total volume of 190 μL. NAAA substrate (heptadecanoyl-ethanolamide, 50 μM) was added and the mixture was incubated at 37°C in a water bath for 2 h. Reactions were stopped with 0.6 mL of stop solution (cold chloroform: methanol; 2:1 v/v) containing 1 nmol heptadecanoic acid (NuChek Prep, Waterville, MN) as internal standard. Samples were centrifuged at 2095 x *g* for 15 min (4°C); organic phases were collected, dried under nitrogen, and resuspended in 75 μL of methanol. Samples (injection volume 5 μL) were eluted isocratically on an Acquity UPLC BEH C18 column (50 mm length, 2.1 mm i.d., 1.7 mm pore size, Waters, Milford, MA) at 0.5 mL/min for 1.5 min with a solvent mixture of 95% methanol and 5% water, both containing 0.25% acetic acid and 5 mM ammonium acetate. Column temperature was set at 40°C. Electrospray ionization was in the negative mode, capillary voltage was 2.7 kV, cone voltage was 45 kV, and source temperature was 150°C with a desolvation temperature of 450°C. Nitrogen was used as drying gas at a flow rate of 800 L/h and a temperature of 500°C. The [M-H]^-^ ion was monitored in the selected-ion monitoring mode *(m/z* values: heptadecenoic acid 267.37, heptadecanoic acid, 269.37). Calibration curves were generated using authentic heptadecenoic acid.

### Real-Time quantitative PCR

Total RNA was prepared from tissue samples (3-5 mg) or cell pellets (2.5×10^5^ cells) using the Ambion PureLink RNA minikit (Life Technologies, Carlsbad, CA, USA) as directed by the supplier. Samples were treated with DNase (PureLink DNase, Life Technologies) and cDNA synthesis was carried out using the Super-Script VILO cDNA synthesis kit (Life Technologies) according to the manufacturer’s protocol using purified RNA (1 μg). *Cell extracts.* First-strand cDNA of SHSY5Y cells was amplified using the iQ SYBR Green SuperMix (Life Technologies) according to the manufacturer’s instructions. Primers were purchased from Origene Technologies (Rockville, MD, USA). Quantitative PCR was performed in a 96-well PCR plate and run at 95°C for 10 min, followed by 40 cycles, each cycle consisting of 15 s at 95°C and 1 min at 60°C, using a ViiA7 instrument (ViiATM 7 real-time PCR system, Life Technologies). The sequences of primer for human *Naaa* are: forward sequence: ATTACGACCACTGGAAGCCAGC; reverse sequence: GGAAAAGTGCCTCCAGGCTGAG. The BestKeeper software (Pfaffl *et al*, 2004) was used to determine expression stability and geometric mean of 4 different housekeeping genes (GAPDH, 18S, ACTB, and HPRT). ΔCt values were calculated by subtracting the Ct value of the geometric mean of these housekeeping genes from the Ct value for the genes of interest. The relative quantity of genes of interest was calculated by the expression 2-ΔΔCt.

#### Tissue extracts

Total RNA was extracted and retro-transcribed from SN, dorsalateral striatum and human cortex as described above. Quantification was performed by a sequence detector (ViiATM 7 Real-Time PCR System; Applied Biosystems, USA) using the TaqMan 5’ nuclease activity from the TaqMan Universal PCR Master Mix, fluorogenic probes, and oligonucleotide primers. Copy numbers of cDNA targets were quantified by the point during cycling when the PCR product was first detected. Gene-specific primers for Taqman assays were purchased from Life Technologies. TaqMan assays were done in triplicate. The mRNA expression levels for mice tissue samples were normalized to the level of the housekeeping gene Hprt. Human samples were normalized on the geometric mean of Actin, Cytochrome C1 and GAPDH.

### Proteomic analyses

*Sample preparation.* Midbrain tissue fragments containing the left or right SN were dissected from 3 independent biological replicates and homogenized in RIPA buffer. Protein concentrations were measured using the BCA assay, and equal amounts of protein were collected and transferred to Eppendorf tubes. Samples were isotopically labeled using TMT sixplex kits (Rauniyar *et al*, 2013), according to the manufacturer’s instruction. The following labeling scheme was used:

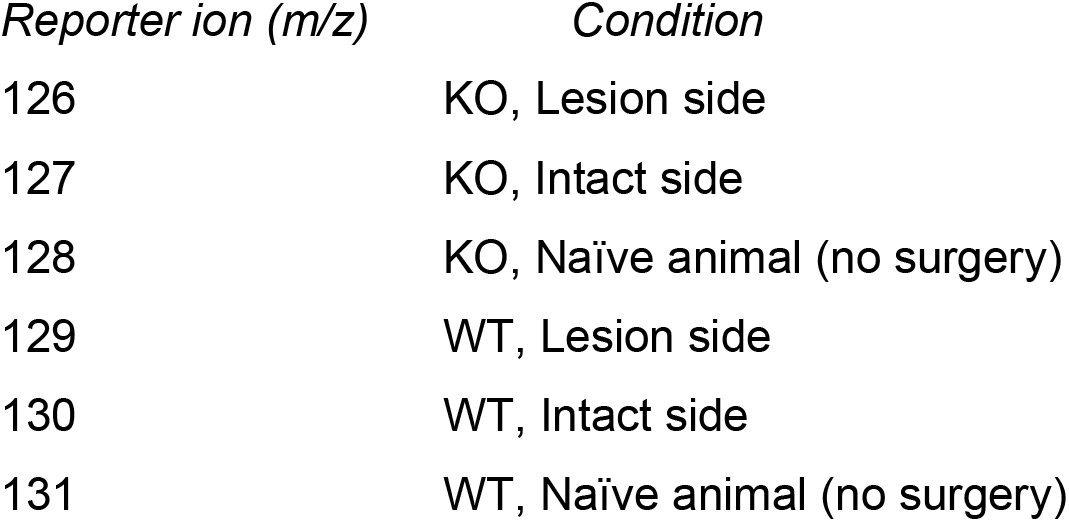

| *Reporter ion (m/z)* | *Condition* |
| --- | --- |
| 126 | KO, Lesion side |
| 127 | KO, Intact side |
| 128 | KO, Naïve animal (no surgery) |
| 129 | WT, Lesion side |
| 130 | WT, Intact side |
| 131 | WT, Naïve animal (no surgery) |

Samples were reduced with dithiothreitol, alkylated with iodoacetic acid and labeled using TMT tags. After pooling the six conditions, the total protein content from each replicate (150 μg) was separated on precast polyacrylamide gels (Thermo Fisher Scientific). Each lane was cut into 12 slices and in-gel protein digestion was performed (Shevchenko *et al*, 2007). The samples were recovered, dried under N2, and dissolved in 50 μL of water containing acetonitrile (3%) and formic acid (0.1%) for LC-MS/MS analysis. *LC-MS/MS analyses.* Tryptic peptide mixtures were analyzed using a Synapt G2 QToF equipped with a nanoACQUITY liquid chromatographer and a nanoSpray ion source. Peptides were separated on a BEH nanobore column (75 μm ID x 25 cm length) using a linear gradient of acetonitrile in water from 3% to 55% in 3 h, followed by a washout step with 90% acetonitrile (10 min) and a reconditioning step in 3% acetonitrile for 20 min. Flow rate was set to 300 nL/min. Spray voltage was 1.6kV, cone voltage was 28V, spray gas was 0.3 L/min. Survey spectra were acquired over the *m/z* = 50-1600 scan range. Multiply charged ions (2+,3+ and 4+) between *m/z* = 300 and 1400 were selected as precursors for DDA tandem mass analysis and fragmented in the trap region of the instrument. Collision energy values were automatically selected by the software using dedicated charge-state dependent CE/*m/z* profiles. Every 60s, a single leu-enkefalin (2 ng/mL) MS scan was acquired by the LockMass ion source for spectra recalibration. *Data analysis.* Acquired raw data files were processed using PLGS software to recalibrate the mass spectra and generate the precursor-fragments peak list. Protein identification and quantification was performed by interrogating the SwissProt database using MASCOT Server software. The search parameters were set as follows, quantification: TMT6plex; fixed modifications: carbamidomethyl (C), TMT6plex (N-term), TMT6plex (K); variable modifications: acetyl (K), acetyl (N-term), deamidated (NQ), methyl (DE), oxidation (M), phospho (ST), phospho (Y); peptide tolerance: 30ppm; fragment tolerance: 0.5Da; maximum allowed missed cleavages: two. At least two peptides were required for a positive protein identification and quantification. A significance threshold of P>0.05 was set for protein identification in the MASCOT search. All the proteins identified and quantified from all the biological replicates were retained. Proteins sharing the same set or subset of peptides were considered as a single entry (MASCOT Protein Families). Two different data analyses were then performed on the same data set. We first investigated the alterations caused by 6OHDA on the brain of naïve animals. For this purpose, only the ratio between 129 and 130 reporter ions was calculated and reported. Protein expression ratio was normalized by the average ratio of all the peptides assigned to proteins. A standard deviation of the protein expression ratios was calculated for all the proteins quantified in at least two out of three replicates. All the corresponding data are available in Supplementary Table 3. We then evaluated the role of the genotype (NAAA^-/-^ versus WT) on the expression profiles of proteins in the three biological conditions: naïve animal, lesioned and contralateral side of the brain. For this purpose, three ratios were measured using the 6plex TMT reporter ions and the reporting function of MASCOT: 126/129 for the lesion side, 127/130 for the contralateral side and 128/131 for the naïve brain. Protein expression ratio was normalized by the average ratio of all the peptides assigned to proteins. Protein expression ratio was normalized by the average ratio of all the peptides assigned to proteins. A standard deviation of the protein expression ratios was calculated for all the proteins quantified in at least 2 out of three replicates. All the corresponding data are available in Supplementary Table 9.

### Neurotransmitter measurements

*6-OHDA experiments:* Striata were removed, snap frozen in liquid nitrogen and stored at −80°C. Samples were weighed and homogenized in 0.9 mL/methanol:water (1:1) containing 0.1% formic acid. After stirring and centrifugation, the supernatants were dried under nitrogen and the samples were reconstituted in 90 μL of mobile phase A (water containing 0.1% acetic acid) for LC-MS/MS analyses. LC-MS/MS analyses were carried out on an Acquity UPLC system coupled with a Xevo TQ-MS triple quadrupole mass spectrometer. Chromatographic separation was achieved using a BEH C18 column (2.1×100 mm, 1.7 micron particle size) eluted at a flow rate of 0.35 mL/min, using the following gradient: 0-1.0 min 5% solvent B (methanol) in solvent A (0.1 % acetic acid in water), 1.0-2.5 min 5% to 100% B, and 2.5-3.5 min 100% B. The column was re-equilibrated to initial conditions from 3.5 to 4.5 min. Total run time for analysis was 4.5 min, and injection volume was 5 μL. The column temperature was kept at 45°C. The MS was operated in both positive and negative ESI mode with cone voltage, collision energy and capillary voltage set at 10V, 20V and 3kv respectively. The source temperature was set to 120°C. Desolvation gas and cone gas (nitrogen) flow were set to 800 and 50 L/h, respectively. Desolvation temperature was set to 450°C. Analytes were quantified by multiple reaction monitoring (MRM) with the following transitions (*m/z*): dopamine, 153.8>136.7; serotonin 176.9>159.9; DOPAC, 166.5>122.9; and HVA, 180.7>136.6. Dopamine and serotonin were acquired in positive mode and DOPAC and HVA in negative mode. Data were acquired by MassLynx software and quantified by TargetLynx software; a linear standard calibration curve was used for analyte quantification.

#### MPTP experiments

A stock solution (1 mg/mL) of HVA, DOPAC, DA and serotonin was prepared in 100 mM sodium borate buffer. The stock solution (10 μL) was mixed with [^2^H_5_]-benzoyl chloride (10 μL) in 100 mM sodium borate buffer and vortexed for 30 s. The mixture was incubated at room temperature for 30 min. The precipitate was dissolved in acetonitrile and methanol was added to quench the reaction, yielding a final concentration of 10 μg/mL internal standard (ISTD). The dorsal striatum (~15 mg) was homogenized in 0.5 mL ice-cold acetonitrile containing 1% formic acid and 10 μg/mL ISTD. The samples were stirred vigorously for 30 s and centrifuged at 2800 × *g* at 4°C for 15 min. After centrifugation, the supernatants were transferred to 5 mL glass vials. Tissue pellets were rinsed with water/acetonitrile (1:4, v/v; 0.2 mL), stirred for 30 s, and centrifuged at 2800 × *g* at 4°C for 15 min. The supernatants were transferred and pooled with the first eluate. Eluates were dried under N_2_ and reconstituted in 0.1 mL of 100 mM Na_2_BO_4_ and vortexed. Benzoyl Chloride (5 μL) was added and the mixture was vortexed and allowed to incubate at room temperature for 30 min. After incubation, methanol (50 μL) was added to quench the reaction, followed by acetonitrile to dissolve the resulting precipitate. Samples were filtered through Agilent Captiva syringe filters in to deactivated glass inserts (0.2 mL) placed inside amber glass vials (2 mL; Agilent Technologies). LC separations were carried out using a 1200 series LC system (Agilent Technologies), consisting of a binary pump, degasser, thermostated autosampler and column compartment coupled to a 6410B triple quadrupole mass spectrometric detector (MSD; Agilent Technologies). Analytes were separated on an Eclipse PAH C18 column (1.8 μm, 2.1 × 50.0 mm; Agilent Technologies). The mobile phase consisted of water containing 0.1% formic acid and 5 mM ammonium formate as solvent A and methanol containing 0.1% formic acid and 5 mM ammonium formate as solvent B. The flow rate was 0.4 mL/min. The gradient conditions were as follows: starting 40% B to 62% B in 7.0 min, changed to 95% B at 7.01 min and maintained till 8.0 min, changed to 40% B at 8.01 min and maintained until 10.0 min for re-equilibration. The column temperature was maintained at 30°C and the autosampler at 9°C. The total analysis time, including re-equilibrium, was 10 min. The injection volume was 2 μL. To prevent carry over, the needle was washed in the autosampler port for 30 s before each injection using a wash solution consisting of 10% acetone in water/methanol/isopropanol/acetonitrile (1:1:1:1, v/v). The MS was operated in the positive electrospray ionization (ESI) mode, and analytes were quantified by multiple reaction monitoring (MRM) of the following transitions: benzoyl-DA 466.2 > 77.1 *m/z,* [^2^H_5_]-Benzoyl-DA 481.3 > 110.1 *m/z,* benzoyl-DOPAC 394.1 > 77.1 *m/z,* [^2^H_5_]-benzoyl-DOPAC 404.2 > 110.1 *m/z,* benzoyl-SE 385.2 > 77.1 *m/z,* [^2^H_5_]-Benzoyl-SE 395.2 > 110.1 *m/z,* benzoyl-HVA 304.1 > 77.1 *m/z,* [^2^H_5_]-benzoyl-HVA 309.2 > 110.1 *m/z.* The capillary voltage was set at 2500 V. The source temperature was 350°C and gas flow was set at 10.0 L/min. Nebulizer pressure was set at 45 psi. Collision energy and fragmentation voltage were set for each analyte as reported. The MassHunter software (Agilent Technologies) was used for instrument control, data acquisition, and data analysis.

### Apomorphine-induced rotations

Three weeks after 6-OHDA administration groups of 15 to 20 mice received subcutaneous injections of apomorphine (dissolved in saline containing 0.2 mg/mL ascorbic acid) and motor behavior was assessed for the following 60 min. Rotational asymmetry was assessed using the ANY-maze Behavior Tracking Software (Stoelting Europe, Dublin, Ireland). Only full body turns were counted (Ungerstedt & Arbuthnott, 1970). Data are presented as total number of net turns in 1 h, with rotation toward the side contralateral to the lesion given a positive value.

### Rotarod test

Behavioral testing on a mouse rotarod apparatus equipped with a 3 cm diameter rod (Ugo Basile, Varese, Italy) was performed on day 21 after surgery, or 7 days post MPTP intoxication, with an accelerating rod (from 15 to 25 rpm in 150 s). All mice were pre-trained one week before the treatments. The training consisted of 5 consecutive runs at 20 rpm or until the mice were able to maintain themselves for 300 s on the rotating rod. Subsequently, a new test series with accelerating conditions was started. One day prior to surgery, baseline values were obtained under the same conditions.

### Statistical analyses

Data were analyzed as described in figure legends using GraphPad Prism version 5 for Windows (La Jolla, California, USA). For data with Gaussian distribution, parametric statistical analysis was performed using the two-tailed Student’s t-test for two groups; one-way or two-way analysis of variance (ANOVA) was applied for multiple comparisons with Bonferroni post hoc analysis for data meeting homogeneity of variance.

## ACKNOWLEDGMENTS

The work was partially funded by grant R01AG065329 (to DP and KG). The technical assistance of Andrea Armirotti is gratefully acknowledged. We are also grateful to the Banner Sun Health Research Institute Brain and Body Donation Program of Sun City, Arizona for the provision of human brain tissue. The Brain and Body Donation Program is supported by the National Institute of Neurological Disorders and Stroke (U24 NS072026 National Brain and Tissue Resource for Parkinson’s Disease and Related Disorders), the National Institute on Aging (P30 AG19610 Arizona Alzheimer’s Disease Core Center), the Arizona Department of Health Services (contact 211002, Arizona Alzheimer’s Research Center), the Arizona Biomedical Research Commission (contracts 4001, 0011, 05-901 and 1001 to the Arizona Parkinson’s Disease Consortium) and the Michael J. Fox Foundation for Parkinson’s Research.

## AUTHOR CONTRIBUTION

S. Pontis designed and performed 6-OHDA experiments, participated in the writing of the manuscript. F. Palese designed and performed experiments on human samples and MPTP investigations and participated in the writing of the manuscript. N. Realini performed experiments on cell lines and lipid measurements. A. Torrens and F. Ahmed performed neurotransmitter quantification experiments. F. Assogna, C. Pellicano and P. Bossù were involved in the recruitment of patients and blood collection. G. Spalletta participated in the interpretation of human data and in the writing of the manuscript. K. Green participated in the writing of the manuscript. D. Piomelli ideated and supervised the study and wrote the manuscript.

## CONFLICT OF INTEREST

The authors declare the following conflict of interest: DP is an inventor in patents issued to the University of California, the University of Parma, and the University of Urbino ‘Carlo Bo’, which protect composition of matter and uses of NAAA inhibitors.

## THE PAPER EXPLAINED

PROBLEM: Available treatments for Parkinson’s disease address the symptoms but not the cause of this debilitating disorder in which dopamine-releasing neurons of the brain undergo progressive degeneration. To overcome this impasse, we need to identity molecular events that control dopamine neuron degeneration and can be targeted by disease-modifying therapies.

RESULTS: We hypothesized that an enzyme called NAAA might control dopamine neuron survival in PD. We tested this idea in three ways. First, we measured NAAA levels in samples from persons with PD and found them to be higher than in persons without PD. Second, we carried out similar measurements in human neurons in cultures or in live mice treated with two toxins that selectively damage dopamine neurons. In both experiments, the toxins strongly increased NAAA levels in neurons. Finally, we asked whether removing the NAAA gene or administering a compound that selectively blocks NAAA activity would alter dopamine neuron degeneration and its behavioral consequences in mice treated with the toxins. Both interventions strongly protected the animals.

IMPACT: Drugs that inhibit NAAA activity might find application as disease-modifying agents that counter the root cause of PD.

## DATA AVAILABILITY

This study includes no data deposited in external repositories.

## SUPPLEMENTARY FIGURES

**Supplementary Figure 1. NAAA is expressed by neurons and microglial cells in postmortem cortex samples from subjects with PD.**

• (A-L) Immunofluorescence images of postmortem cortex samples from subjects with. Scale bar, 20 μm. NAAA is shown in green (B, F, J), Map2 in magenta (C, D), Iba1 in red (G, H), GFAP in grey (K, L). Cell nuclei are stained with DAPI (blue, A, E, I).

**Supplementary Figure 2. 6-OHDA stimulates early NAAA expression in nigral dopamine neurons.**

• (A-D1) Immunofluorescence images of tissue sections prepared 48 h after 6-OHDA injection from (A-D) control SN (contralateral to 6-OHDA injection site) and (A1-D1) lesioned SN (ipsilateral to 6-OHDA injection site). Scale bar, 50 μm.

• (E-I1) High-magnification images of sections from (E-I) control SN and (E1-I1) lesioned SN. Scale bar, 10 μm. NAAA is shown in green, TH in red, Iba-1 in magenta. Merged signals are shown in yellow. Cell nuclei are stained with DAPI (blue).

**Supplementary Figure 3. 6-OHDA stimulates delayed NAAA expression in nigral microglia.**

• (A-D1) Immunofluorescence images of tissue sections prepared 14 days after 6-OHDA injection from (A-D) control SN (contralateral to 6-OHDA injection site) and (A1-D1) lesioned SN (ipsilateral to 6-OHDA injection site). Scale bar, 50 μm.

• (E-I1) High-magnification images of sections from (E-I) control SN and (E1-I1) lesioned SN. Scale bar, 10 μm. NAAA is shown in green, TH in red, Iba-1 in magenta. Merged signals are shown in white. Cell nuclei are stained with DAPI (blue).

**Supplementary Figure 4. Time-course of NAAA expression in the SN**.

• (A-A2) Immunofluorescence images of tissue sections prepared from the SN pars compacta 24h after 6-OHDA injection; (A) TH (red), (A1) NAAA (green) and (A2) merged (yellow).

• (B-B2) SN images 48 h after 6-OHDA injection: (B) TH (red), (B1) NAAA (green) and (B2) merged (yellow).

• (C-C2) confocal images of SN 7 days after 6-OHDA: (C) TH (red), (C1) NAAA (green) and (C2) merged (yellow).

• (D) Quantitative co-localization analysis: co-localization values are expressed as mean ± SD (n=6) (for details, see Online Methods). For each image, the Costes’ P-value from Costes’ randomized analysis was 100%, indicating maximum probability that the colocalization was not due to chance.

**Supplementary Figure 5. 6-OHDA causes rapid proteome-wide alterations in the SN.**

• Comparative proteomics analyses of SN tissue identified and quantified 1164 proteins *(Table S1),* 417 of which were differentially expressed by more than 20% between lesioned and intact SN in at least two of the three biological replicates included in our analyses. The majority of deregulated proteins (392) were underrepresented. Annotation of underrepresented proteins using Gene Ontology (GO) (http://www.geneontology.org) tools shows enrichment in proteins that clustered in the following categories: (A) cellular components; (B) molecular function; and (C) biological process.

**Supplementary Figure 6. Effect of 6-OHDA on oxidative phosphorylation, carbon metabolism and lipid and glycan metabolism pathways.**

• Comparative proteomics analysis of SN tissue (intact versus lesioned: ipsilateral and contralateral, respectively, to the 6-OHDA injection site): (A) oxidative phosphorylation; (B) carbon metabolism; and (C) lipid and glycan metabolism.

**Supplementary Figure 7. 6-OHDA stimulates delayed NAAA expression in striatal microglia.**

• (A-E) Immunofluorescence images of tissue sections from dorsolateral striatum prepared

14 days after 6-OHDA injection. Cell nuclei (DAPI, blue); NAAA (green), TH (red), Iba-1 (magenta) and merged (yellow). Immunofluorescence images from intact (B-E) and lesioned sides (B1-E1) highlighting (B, B1) increased NAAA levels; (C, C1) loss of TH^+^ fibers; and (D1, D) increased Iba-1 immunofluorescence. Scale bar, 50 μm.

• (F-I1) High-magnification images of sections stained with DAPI (F, F1, blue), NAAA (G, G1, green), Iba-1 (H, H1, magenta) and merged (i, i1, yellow). Scale bar, 20 μm. Images from lesioned striatum show that NAAA^+^/Iba-1^+^ cells have the typical morphology of reactive microglia (G-I1).

**Supplementary Figure 8. Biochemical characterization of *Naaa^-/-^* and *Naaa^+/-^* mice.**

• (A-B) RT-PCR quantification of NAAA mRNA levels in (A) brain and (B) lungs; wild-type mice, WT. ***P<0.001, **P<0.01, one-way ANOVA followed by Bonferroni post hoc test (n=4).

• (C) Densitometric quantification (C1) and representative western blot (C2) of NAAA levels in mouse brain homogenates; NAAA signal was normalized to GAPDH. **P<0.01 Student’s *t* test (n=4).

• (D) NAAA activity in lung tissue extracts. **P<0.01, one-way ANOVA followed by Bonferroni post hoc test (n=4).

• (E-H) RT-PCR quantification of Fatty acid amide hydrolase *(FAAH, E),* N-acylphosphatidylethanolamine phospholipase D *(NAPE-PLD,* F), Acid ceramidase *(ASAH1,* G) and ß-glucocerebrosidase-1 *(GBA,* H) mRNA levels in brain (n=6).

**Supplementary Figure 9. Effects of 6-OHDA on striatal content of dopamine metabolites and serotonin in Naaa^-/-^ mice.**

• (A-C) Levels of (A) 3,4-dihydroxyphenylacetic acid (DOPAC), (B) homovanillic acid (HVA), and (C) serotonin (5-HT) in striatal extracts prepared three weeks after 6-OHDA injection. *P<0.05, one-way ANOVA followed by Bonferroni post hoc test (n=4).

**Supplementary Figure 10. Effects of 6-OHDA on striatal content of dopamine metabolites and serotonin in mice treated with ARN19702.**

• (A-C) Levels of (A) 3,4-dihydroxyphenylacetic acid (DOPAC), (B) homovanillic acid (HVA), and (C) serotonin (5-HT) in striatal extracts prepared 3 weeks after 6-OHDA injection (n=4).

**Supplementary Figure 11. Effects of MPTP on striatal content of dopamine metabolites and serotonin in *Naaa^-/-^* mice.**

• (A-C) Levels of (A) 3,4-dihydroxyphenylacetic acid (DOPAC), (B) homovanillic acid (HVA), and (C) serotonin (5-HT) in striatal extracts prepared 3 weeks after MPTP injection (n=3-4).

**Supplementary Figure 12. Effects of MPTP on striatal content of dopamine metabolites and serotonin in mice treated with ARN19702.**

• (A-C) Levels of (A) 3,4-dihydroxyphenylacetic acid (DOPAC), (B) homovanillic acid (HVA), and (C) serotonin (5-HT) in striatal extracts prepared 3 weeks after MPTP injection (n=3-5).

**Supplementary Figure 13. Genetic or pharmacological NAAA blockade reduces dopaminergic neuronal loss in MPTP-treated mice.**

• (A-E) Representative immunofluorescence images of dopaminergic (TH+) neurons in SN 7 days after MPTP (B, D, F,) or saline administration (control, A, C,E) in vehicle (A,B), ARN19702 (C,D) treated mice or *Naaa^-/-^* mice (E,F). Scale bar, 100 μm. TH is shown in green, cell nuclei are stained with DAPI (blue).

**Supplementary Figure 14. Genetic or pharmacological NAAA blockade reduces dopaminergic fibers degeneration in MPTP-treated mice.**

• (A-F) Representative immunofluorescence images of dopaminergic (TH+) fibers in dorsal striatum 7 days after MPTP (B, D, F,) or saline administration (control, A, C,E) in vehicle (A,B), ARN19702 (C,D) treated mice or Naaa^-/-^ mice (E,F). Scale bar, 100 μm. TH is shown in magenta, cell nuclei are stained with DAPI (blue).

**Supplementary Figure 15. Comparative proteomics analysis of intact and lesioned SN from wild-type and *Naaa^-/-^* mice.**

• (A-C) Analyses of lesioned SN tissue (ipsilateral to the 6-OHDA injection site) identified 1164 quantifiable proteins (Supplementary *Table 7),* 619 of which were differentially expressed between wild-type and NAAA^-/-^ mice (Supplementary *Table 8).* Gene Ontology (GO) tools (http://www.geneontology.org) assigned overrepresented proteins to one of the following categories: (A) biological process; (B) molecular function; (C) cellular component.

## SUPPLEMENTARY TABLES

**Supplementary Table 1.** Sociodemographic and clinical characteristics of subjects included in the present study: postmortem analyses. Race, 1: white. Biological sex, 1: male; 2: female. PMI: Postmortem interval.

**Supplementary Table 2.** Sociodemographic and clinical characteristics of subjects included in the present study: premortem analyses. H&Y: Hoehn and Yahr score; MMSE: Mini Mental State Examination. UPDRS III: Unified Parkinson’s Disease Rating Scale, Motor Examination.

**Supplementary Table 3.** Comparative proteomics analysis of SN tissue 48 h after 6-OHDA injection: intact versus lesioned side (lesioned: ipsilateral to 6-OHDA injection site; intact: contralateral to 6-OHDA injection site) from wild-type mice. The table lists SwissProt accession number and annotation for all quantifiable proteins identified in the three biological replicates included in the present study.

**Supplementary Table 4.** Proteins differentially expressed in lesioned versus intact SN tissue of wild-type mice (lesioned: ipsilateral to 6-OHDA injection site; intact: contralateral to 6-OHDA injection site). The 417 proteins included in the list (of a total of 1194 quantifiable proteins) are those present in at least two of the three biological replicates included in the present study. Overexpressed: blue background; underexpressed: green background.

**Supplementary Table 5**: Select protein pathways differentially expressed in lesioned versus intact SN tissue of wild-type mice (lesioned: ipsilateral to 6-OHDA injection site; intact: contralateral to 6-OHDA injection site). The list includes proteins that were detected in at least two of the three biological replicates included in the present study. KEGG: Kyoto Encyclopedia of Genes and Genomes.

**Supplementary Table 6:** Partial list of proteins in the oxidative phosphorylation pathway, which are differentially expressed in lesioned versus intact SN tissue of wild-type mice (lesioned: ipsilateral to 6-OHDA injection site; intact: contralateral to 6-OHDA injection site).

**Supplementary Table 7:** Partial list of proteins in carbon metabolism pathways, which are differentially expressed in lesioned versus intact SN tissue (lesioned: ipsilateral to 6-OHDA injection site; intact: contralateral to 6-OHDA injection site).

**Supplementary Table 8:** Partial list of proteins in lipid and glycan metabolism pathways, which are differentially expressed in lesioned versus intact SN tissue (lesioned: ipsilateral to 6-OHDA injection site; intact: contralateral to 6-OHDA injection site).

**Supplementary Table 9.** Comparative proteomics analysis of SN tissue 48h after 6-OHDA injection: wild-type versus *Naaa^-/-^* mice. The table lists SwissProt accession number and annotation for all quantifiable proteins identified in the three biological replicates included in the present study.

**Supplementary Table 10.** Proteins differentially expressed in lesioned SN tissue from *Naaa^-/-^* versus wild-type mice (lesioned: ipsilateral to 6-OHDA injection site; intact: contralateral to 6-OHDA injection site). The 619 proteins included in the list (of a total of 1194 quantifiable proteins) are those present in at least two of the three biological replicates included in in the present study. Under-expressed: blue background; overexpressed: green background.

**Supplementary Table 11**. Select protein pathways differentially expressed in lesioned SN tissue from *Naaa*^-/-^ versus wild-type mice (lesioned: ipsilateral to 6-OHDA injection site; intact: contralateral to 6-OHDA injection site). The list includes proteins that were detected in at least two of the three biological replicates included in the present study. KEGG: Kyoto Encyclopedia of Genes and Genomes.

**Supplementary Table 12.** Partial list of proteins in the oxidative phosphorylation pathway, which are differentially expressed in lesioned SN tissue of *Naaa*^-/-^ versus wild-type mice (lesioned: ipsilateral to 6-OHDA injection site; intact: contralateral to 6-OHDA injection site).

**Supplementary Table 13.** Partial list of proteins in the carbon metabolism pathways, which are differentially expressed in lesioned SN tissue of *Naaa*^-/-^ versus wild-type mice (lesioned: ipsilateral to 6-OHDA injection site; intact: contralateral to 6-OHDA injection site).

**Supplementary Table 14.** Partial list of proteins in the lipid and glycan metabolism pathways, which are differentially expressed in lesioned SN tissue of *Naaa^-/-^* versus wild-type mice (lesioned: ipsilateral to 6-OHDA injection site; intact: contralateral to 6-OHDA injection site).

**Supplementary Table 15.** PCR conditions for genotyping of *Naaa^-/-^* mice.

**Supplementary Table 16**. Commercial antibodies and experimental conditions utilized in the present study.

